# Chronic Electronic Cigarette Exposure Promotes Atherosclerosis and Chondrogenic Modulation of Smooth Muscle Cells

**DOI:** 10.1101/2025.09.13.675958

**Authors:** Isabella Damiani, Chad S. Weldy, Quanyi Zhao, Elena Hurtado Solberg, Guyu Qin, Meena Easwaran, Siwen Zheng, Sugandha Basu, Wenduo Gu, Matthew Worssam, João Pinho Monteiro, Daniel Li, Gurmenjit Kaur Bahia, Ramendra Kundu, Trieu Nguyen, Elizabeth Erickson-Direnzo, Paul Cheng, Juyong Brian Kim

## Abstract

**Background:** Electronic cigarette (E-cig) use has reached epidemic proportions worldwide, yet its cardiovascular consequences remain poorly defined. While several lines of evidences in human epidemiological and animal studies suggest chronic aerosol exposure accelerates atherosclerosis; the cellular and molecular mechanisms underlying this pathological remain unknown.

**Methods:** We exposed hyperlipidemic mice to chronic e-cigarette aerosol inhalation and characterized the plaque cellular landscape by coupling SMC lineage tracing with single-cell transcriptomic/epigenomic profiling and histologic phenotyping. We subsequently leveraged human coronary artery smooth muscle cells (HCASMCs) to validate *in vivo* discovery and identified E-cig specific pathological signaling pathways relevant to human vascular disease risk.

**Results:** Chronic E-cig aerosol exposure accelerated atherosclerosis, increasing both SMC phenotypic modulation and plaque macrophage burden in a lipid-independent manner. Transcriptomically, SMCs are particularly more sensitive to E-cig than other vascular cell types. E-cig exposure reprogrammed SMCs toward a pro-calcifying, chondrogenic phenotype, thereby enhancing vascular ossification *in vivo* and in HCASMCs *in vitro*. Mechanistically, E-cig mediated SMC fate alteration occurs through activation of a glutamatergic/NMDAR signaling program, that increased NMDAR-dependent Ca^2+^ influx in a *GRIN2A* dependent manner. Notably, inhibition of *GRIN2A* mediated signaling reversed E-cig-induced pathological shifts in SMC phenotype.

**Conclusions:** SMC chondrogenic reprogramming and subsequent vascular calcification are central to the detrimental cardiovascular consequences of E-cig exposure. Our findings implicate a *GRIN2A*-dependent glutamatergic/NMDAR signaling axis in SMC as a primary driver of this calcifying remodeling program. These findings define a SMC-specific vulnerability to E-cig aerosols and establish the *GRIN2A*/NMDAR pathway as potential therapeutic targets for mitigating E-cig-associated cardiovascular disease.

## Introduction

Cardiovascular disease (CVD) remains the leading cause of morbidity and mortality globally, accounting for approximately 20 million deaths each year, underscoring the critical importance of understanding factors influencing its development and progression^1,2^. Traditional tobacco use is a leading risk factor for CVD^3^, and while E-cig was initially promoted as a less harmful alternative to traditional tobacco smoking, emerging evidence suggests that E-cig aerosol exposure may pose substantial risks to vascular health, including the development and progression of atherosclerosis^4–6^.

The global rise in the use of electronic cigarettes (E-cigs) has sparked significant debate regarding their safety and long-term health effects, particularly in relation to CVD^5,7^. Furthermore, E-cig usage has been associated with adverse effects on pulmonary function, increased inflammation, and impaired immune responses, indicating potential risks to multiple organ systems beyond cardiovascular health^8–12^. Although E-cig aerosols generally contain fewer carcinogens than combustible cigarettes, they still deliver toxic aldehydes, volatile organic compounds, and heavy metals that may pose carcinogenic risks with prolonged use^13,14^. These broad systemic effects underscore the need to better understand the mechanisms by which E-cig aerosol exposure contributes to disease risk, particularly in the context of cardiovascular pathologies such as atherosclerosis.

To address the critical knowledge gap regarding the effects of E-cig aerosol exposure on vascular health, this study aimed to investigate its impact on vascular cellular landscape and to elucidate the molecular pathways by which E-cigs accelerate the progression of atherosclerosis. As the most prevalent cell type in the vascular wall^15^, smooth muscle cells (SMCs) play a pivotal role in maintaining vascular integrity and adapting to environmental and pathological stimuli.

In this study, we evaluate the effects of E-cig aerosol exposure on atherosclerosis development using Apolipoprotein E (ApoE) knockout mice with transgenic smooth muscle cell lineage tracing reporter, a well-established mouse model of atherosclerosis. By combining E-cig aerosol exposure with single-cell RNA sequencing (scRNA-seq), single-cell ATAC sequencing (scATAC-seq) analysis and histology, we gain a comprehensive understanding of how different cell types within the atherosclerotic plaque respond to this environmental insult, with a focus on SMC fate.

Our results reveal that E-cig aerosol exposure leads to significant transcriptional and epigenetic changes in the vessel wall, resulting in increased atherosclerosis, and SMC phenotypic modulation, favoring their transition to chondrogenic cell state. Through integration of transcriptomic and epigenomic datasets, we identify glutamate signaling as a potentially novel E-cig induced pathway mediating SMC phenotypic modulation.

Together, our findings provide novel insights into how E-cig aerosol exposure accelerates atherosclerosis, with a particular focus on smooth muscle cell phenotypic modulation, and single-cell multi-omics. The study underscores the need for further research into the cardiovascular risks associated with E-cig use. These insights may inform regulatory policies and interventions aimed at mitigating the public health impact of E-cig aerosol exposure.

## Methods

All data and materials are deposited to the National Center for Biotechnology Information Gene Expression Omnibus (GSE300982, GSE307204). All methods and materials used in this study are described in detail in the Materials and Methods section of the Data Supplement. In the following sections, please see a brief summary of the most relevant details.

### Mouse models

The animal study protocol was approved by the Administrative Panel on Laboratory Animal Care at Stanford University. SMC-specific lineage tracing in the atherosclerotic model was performed as described previously^16^. Tamoxifen was administered by oral gavage at 8 weeks of age to induce the lineage marker, followed by the initiation of a high-fat diet (Dyets No. 101511).

### Murine aortic tissue processing and histology

Immediately after sacrifice, mice were perfused with 4% paraformaldehyde and aortic tissue was harvested, embedded in optimal cutting temperature compound, and sectioned. Immunofluorescence and immunohistochemistry were performed and lesion areas profiled as described previously^16,17^. A full description of staining and quantification methods is provided in the Supplemental Methods.

### In vitro and In vivo exposure

E-cig aerosol (*Juul* Virginia Tobacco flavor) was introduced at 25% v/v into the media for treatment of HCASMC for all *in vitro* experiments including RNA-seq and SMC phenotypic assays. For *in vivo* studies, E-cig aerosol exposures were performed using a whole-body chamber of the inExpose™ system (Scientific Respiratory Equipment Inc [SCIREQ], Montreal, QB, Canada). Mice were exposed to aerosol for 2 hours per day, 3 days per week, for 12 or 16 weeks, starting at 8 weeks of age. *Juul* Virginia Tobacco pods (5.0% nicotine by weight, 59 mg/mL) were selected due to their high nicotine content and widespread use among U.S. adolescents and young adults, ensuring translational relevance.

### Nicotine and Aerosol Chemical Characterization

Aerosol nicotine and other chemical components (propylene glycol and aldehydes including acrolein, formaldehyde, and acetaldehyde) were quantified using selective ion flow tube mass spectrometry (SIFT-MS; Syft Technologies) as described in prior work^18^. Measured nicotine concentrations in the Tedlar bags ranged from 700 to 2000 ppb per 6 puffs, corresponding to an approximate per-puff concentration of 117 to 333 ppb.

### Gravimetric Determination of Total Particulate Matter (TPM)

Total particulate matter (TPM) was determined by weighing filter pads before and after aerosol collection. The average mass increase was 150 mg per pod, yielding a TPM per puff of 1.25 mg/puff (150 mg/120 puffs). This value is consistent with published reports for JUUL devices (1.4 ± 0.4 mg), supporting the validity of our exposure system^19^.

### Analysis of single cell sequencing data

A full description of the methodology for the acquisition and analysis of both single-cell RNA sequencing (scRNA-Seq) and single-cell assay of transposase-accessible chromatin with sequencing (scATAC-Seq) data is provided in the Supplemental Methods.

### *In vitro* genomic and functional studies in primary and immortalized human coronary artery smooth muscle cell (HCASMC)

Primary HCASMC were purchased from Cell Applications, Inc, and experiments were performed with HCASMCs between passages 5 and 8.

### Human Samples

Human coronary artery samples were obtained from the Stanford Department of Cardiothoracic Surgery Human Biorepository Tissue Bank from consenting patients, under protocol approval [Stanford IRB protocol number 67212]. All samples were from heart transplant recipients. Tissue procurement and use were conducted under protocols approved by the Stanford University Institutional Review Board. The basic clinical characteristics of the patients included in this study are presented in Supplementary Table 1 (**Table S1**).

### Statistical methods

R or GraphPad Prism 10.4.2 were used for statistical analysis. Differentially expressed genes in the scRNA-seq and scATAC-seq data were identified using a Wilcoxon rank-sum test. For overlapping of genomic regions or gene sets, we used the Fisher exact test to test for enrichment. For comparison of cell cluster proportions, Chi-square test was used. For comparisons between 2 groups of equal variances, an unpaired 2-tailed t test was performed. For multiple comparisons testing, 1-way ANOVA followed by a Tukey or Dunnett’s post hoc test or 2-way ANOVA followed by a Sidak’s post hoc test were used as appropriate. All error bars represent the mean +/- SEM. Statistical significance is indicated in the graphs as follows: ****P<0.0001, ***P<0.001, **P<0.01, *P<0.05.

## Results

### Long term E-cig aerosol exposure increases atherosclerosis and affects the SMC participation in lesion remodeling

To determine the impact of long term E-cig aerosol exposure on atherosclerotic lesion formation and smooth muscle cell remodeling, we employed the SMC-specific mouse Cre allele (*Myh11-Cre^ERT^*^2^) combined with the floxed tandem dimer tomato (*tdTomato*) fluorescent reporter protein gene knocked into the ROSA26 locus (B6.Cg-Gt(ROSA)26 Sor^tm14(CAG*tdTomato*^/J), in the ApoE^-/-^background (SMC-LnT line). Hyperlipidemia was induced with feeding of a high fat diet (HFD) and the development of atherosclerosis was evaluated with histological analysis of the aortic root. We compared the SMC-LnT mice exposed to E-cig aerosol and HFD (*E-cig*, n=16) to the SMC-LnT mice exposed to Filtered air and HFD (*Ctrl*, n=16) for 16 weeks (**Figure 1A**). To verify systemic nicotine delivery, we quantified urinary nicotine metabolites by LC-MS. Cotinine and trans-3′-hydroxycotinine (3-OH cotinine) were consistently detected in E-cig aerosol-exposed mice but were undetectable in control animals (**Figure 1B,S1A**), confirming effective and reproducible systemic nicotine exposure in our model.

**Figure 1.**
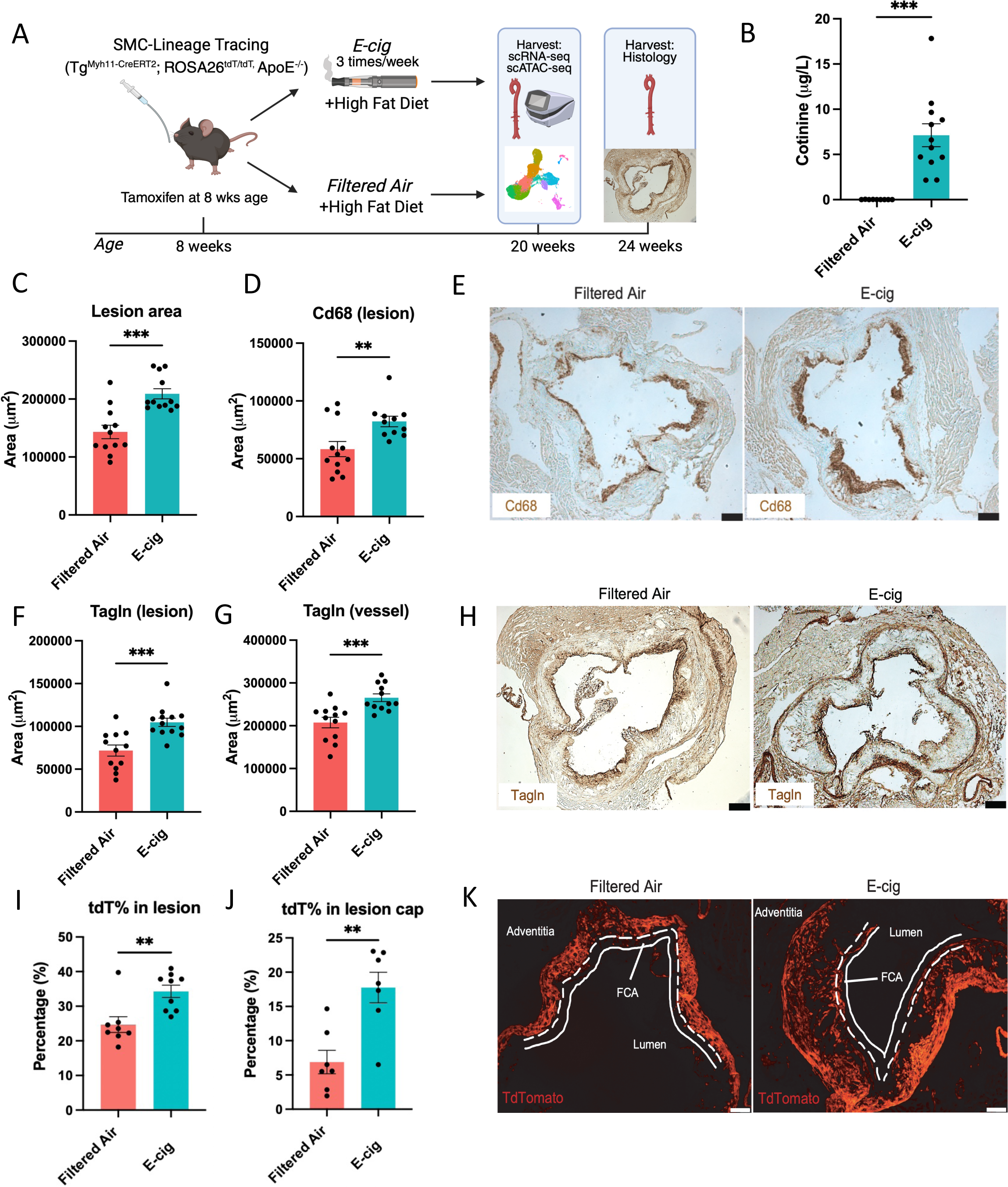
Long term E-cig exposure increases atherosclerosis and affects the SMC participation in lesion remodeling. (A) Schematic figure of E-cig exposure protocol showing SMC-specific lineage tracing mice. (B) Urine cotinine concentrations after 16-week inhalation of E-cig aerosol, P<0.0001. Mice were sacrificed 24h following the last exposure. (C) Total lesion area quantified between the lumen and internal elastic lamina, P= 0.0002. (D) Quantification of Cd68 positive area in the lesion, P=0.0078. (E) Representative aortic root sections from Control and E-cig exposed mice stained for Cd68. Scale bar: 250μm. (F,G) Quantification of Tagln positive SMC areas in lesion and vessel, P=0.0004 in lesion and P=0.0009 in vessel. (H) Representative aortic root sections from Control and E-cig exposed mice stained for Tagln. Scale bar: 250μm.(I,J) Quantification of tdTomato positive area in the lesion and lesion cap, P=0.0043 in the lesion and P=0.0022 in lesion cap. (K) Representative aortic root sections from Control and E-cig exposed mice for tdTomato fluorescence to show SMC lineage-traced cells. Scale bar: 50μm. FCA, fibrous cap area. Data in (B,C,D,F,G,I,J) were analyzed by two-tailed T-test. Data represent mean +/- SEM.

Overall, we found that *E-cig* group had significantly increased total lesion area compared to *Ctrl* group (**Figure 1C**). The Cd68 positive area in the lesions of the E-cig aerosol-exposed mice was also greater compared to Ctrl group suggesting E-cig aerosol exposure results in increased macrophage cell infiltration within the lesion (**Figure 1D, 1E**). To evaluate the involvement of SMC within the plaque we performed immunohistochemical analysis of the SMC-specific marker Transgelin (Tagln). This analysis revealed a significant increase in the overall SMC area within the lesion and vessel following E-cig aerosol exposure compared to Ctrl mice (**Figure 1F-H**). Additionally, we quantified the tdTomato signal to further investigate SMC-lineage cell involvement within the plaque. E-cig aerosol exposure led to a significant increase in the proportion of tdTomato-positive SMC lineage–traced cells in both the lesion and lesion cap, suggesting increased accumulation and/or recruitment of SMC-lineage cells. (**Figure 1I-K**).

To understand if E-cig effect on atherosclerosis development was due to modified lipid profiles, we measured total cholesterol, circulating triglycerides, LDL, and HDL levels and no significant differences were observed between E-cig aerosol-exposed and Ctrl mice **(Figure S1B)**. Consistent with these findings, Oil Red O staining of aortic root lesions did not reveal any statistical difference in lipid deposition between groups **(Figure S1C, S1D)**. Together, these data suggest that E-cig induced effect on atherosclerosis occur independent of changes in circulating lipid levels or intralesional lipid accumulation.

### Long term E-cig aerosol exposure promotes SMC differentiation and modulation to a chondrogenic phenotype

To gain a cellular mechanistic understanding of the effect of E-cig on vascular biology, we exposed SMC-LnT mice for 12 weeks to E-cig or Filtered air with HFD and then isolated the aortic root and ascending aorta and performed single cell multi-omic profiling. Following cellular digestion and mechanical dissociation, we performed FACS sorting for live cells and collected SMC derived (tdTomato-positive) and non-SMC derived (tdTomato-negative) cells. A portion of the single cell suspensions were loaded immediately onto a 10X platform for scRNAseq capture while the remaining cells underwent nuclear isolation prior to 10X capture for scATACseq (see methods). A total of 11 RNA captures (6 captures Filtered air, 5 captures E-cig aerosol-exposed) with 3 mice per capture (total 15-18 mice per exposure group) were completed and subsequent cDNA libraries were sequenced.

Data analysis following our previously reported methods using Seurat with UMAP visualization^20–23^ identified major vascular cell types including endothelial cells (Endo), smooth muscle cells (SMCs), macrophages (MAC) and fibroblasts (Fibro) (**Figure 2A**). Upon analysis of the effect of E-cig aerosol exposure, UMAP visualization revealed a significant transcriptional shift, most notable in the tdTomato-positive SMC population (**Figure 2B, S2A**). To determine the cell type specific transcriptional effect of E-cig aerosol exposure on the vessel wall, we performed differential expression (DE) analysis across SMC, Fibroblast, Macrophage, and Endothelial cells. All four cell types exhibited some degree of transcriptional response, but the most extensive changes were observed in SMCs, followed by fibroblasts, with more modest alterations in macrophages and endothelial cells (**Figure 2D, S2B**). DE profiling in the SMC cluster (**Figure 2C and D**) revealed 638 differentially expressed genes (228 upregulated and 367 downregulated) in response to E-cig, as compared to 323 in fibroblasts, 319 in endothelial cells, and 434 in macrophage cells with overall less fold change effect (**Figure S2B**). These findings indicate that SMCs exhibit the most robust transcriptional response to E-cig aerosol exposure among major vascular cell populations, with a greater number of DE genes and larger effect sizes than fibroblasts, endothelial cells, and macrophages.

**Figure 2.**
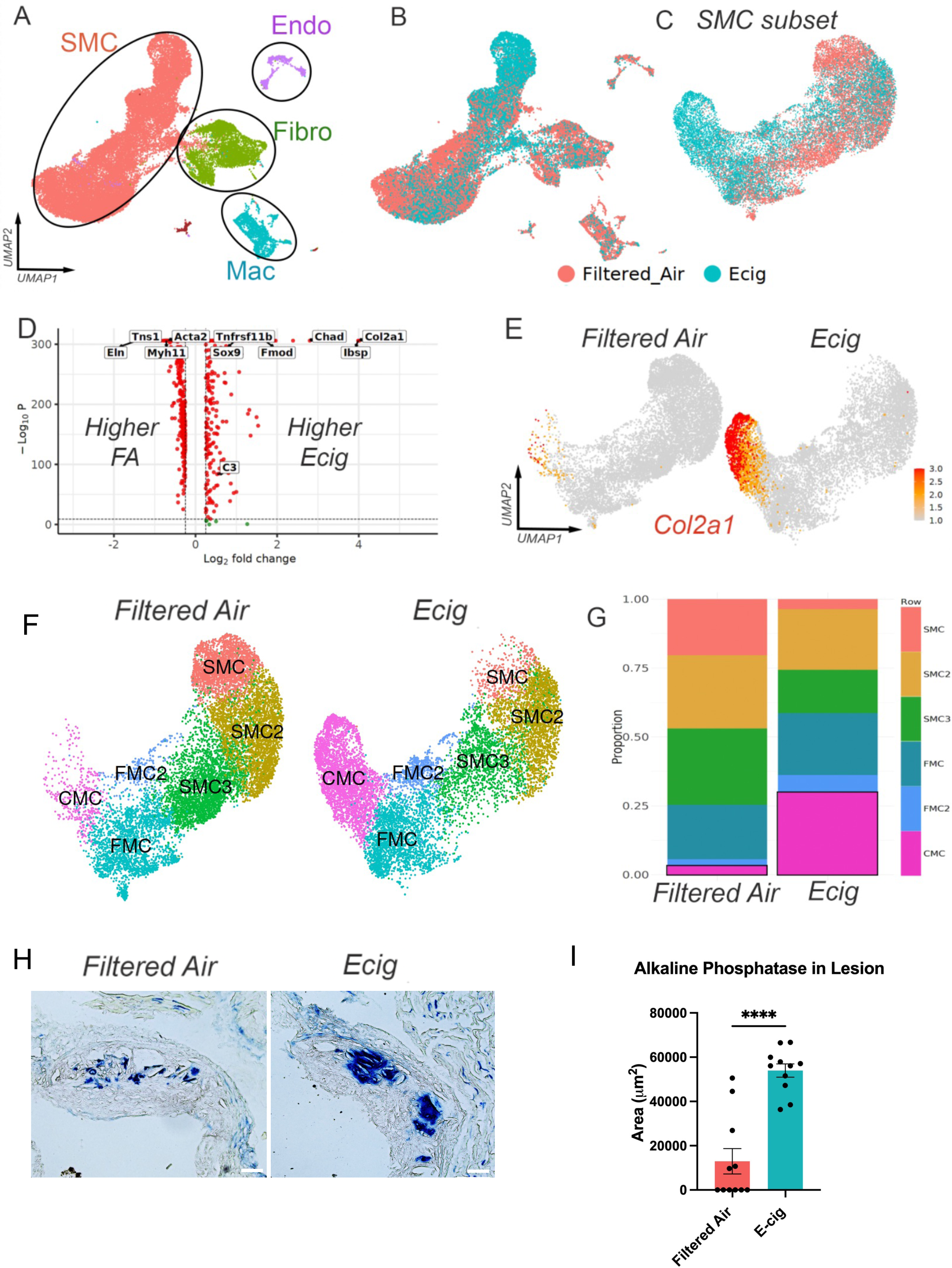
Long term E-cig exposure promotes SMC differentiation and modulation to a chondrogenic phenotype. UMAPs of scRNAseq data from atherosclerotic aortic root and ascending aorta at 12 weeks high fat diet with E-cigarette exposure in SMC lineage tracing mice (SMC-LnT: *Myh11^CreERT^*^2^, *ROSAtdTomato*, *ApoE^-/-^*) showing (A) major cell type clusters (SMC, Fibro, Endo, Mac), (B) grouped by exposure type, and (C) UMAP of SMC subset grouped by exposure type. (D) Volcano plot showing differential gene expression analysis within SMC subset by exposure type. (E) Feature plots of *Col2a1* expression split by exposure type within SMC subset. (F) UMAP clustering of SMC subset split by exposure type. (G) Stacked bar charts showing proportion of cells within SMC clusters split by exposure. (H) Representative images of Ferangi Blue (alkaline phosphatase) calcification staining at the aortic root of SMC-LnT mice following 16 weeks high fat diet grouped by exposure and (I) quantification of Alkaline Phosphatase area, P<0.0001. Scale bar: 50μm (10X).

Among the top upregulated DE genes in SMCs were chondrogenic markers such *as Chad, Col2a1, Sox9, Tnfrsf11b, and Ibsp*, whereas key SMC contractile markers (*Acta2, Myh11*) and cell adhesion genes (*Eln, Tns1*) were among the most downregulated (**Figure 2D**). Feature plot analysis of *Col2a1*, a chondromyocyte marker, revealed striking upregulation in SMCs following E-cig aerosol exposure (**Figure 2E**). To further investigate the functional consequences of these transcriptomic shifts, we performed pathway enrichment analysis of DE genes in SMCs, which revealed upregulation of ossification and biomineralization pathways and downregulation of muscle cell differentiation and development programs (**Figure S2C**). Together, these results indicate that E-cig aerosol exposure promotes a phenotypic shift in SMCs toward a chondrogenic and mineralizing state, potentially contributing to plaque vulnerability and vascular calcification.

Further unbiased clustering analysis of the SMC subset revealed 6 main clusters that were annotated as SMC, SMC2, SMC3, fibroblast-like ‘fibromyocytes’ (FMCs), pro-inflammatory FMCs (FMC2) and chondrogenic ‘chondromyocytes’ (CMCs), as previously described (**Figure 2F**).^20,23,24^ SMC, SMC2, and SMC3 clusters were characterized by expression of mature SMC markers, including *Cnn1* and *Myh11*. In contrast, FMCs were defined by downregulation of mature SMC markers and upregulation of genes such as *Vcam1* and *Fbln2* (**Figure S2D**). FMC2 exhibited high expression of *Fbln1*, *Cxcl12*, and *C3*, whereas CMCs were marked by *Col2a1* and *Spp1* expression (**Figure S2D**).

Following E-cig aerosol exposure, we found a significant shift of the SMC lineage cells toward a CMC phenotype. The proportion of CMCs was markedly higher in the E-cig group compared to the Ctrl group (3.2% in control versus 30.0% in E-cig, *p<E-16*). This shift was accompanied by a reduced proportion of mature SMCs (SMC/SMC2/SMC3) and an increase in the FMC2 population (**Figures 2F,G**). Our previous results have found that increased CMC content is associated with increased vascular calcification^20,24^. Subsequent histological analysis of plaque area through measurement of alkaline phosphatase activity (AP) further confirmed E-cig to increase vascular ossification activity (**Figures 2H,I**). Overall, these findings show that E-cig aerosol exposure can significantly impact SMC phenotype and drive SMC towards a pro-calcifying CMC phenotype, contributing to vascular calcification in the context of atherosclerosis.

### E-cig activates a cluster of pro-inflammatory SMC

In addition to increasing CMC formation, E-cig increased the proportion of SMC-lineage cells within the FMC2 cluster that lies at the intersection between mature contractile SMC clusters (SMC/SMC2/SMC3) and the CMC cluster (2.4% in Ctrl versus 6.1% in E-cig, *p<E-16*) (**Figure 2G**). This FMC2 cluster was marked by proinflammatory markers *C3*, *Cxcl12*, and *C4b* with decreased expression of SMC contractile markers *Myh11* and *Acta2* (**Figure S2D**). To determine the spatial distribution of this cell population within atherosclerotic plaques, we performed RNAscope analysis for *C3*, *Cxcl12* and *Fbln1* in sections of the atherosclerotic lesions (**Figure S2E**). RNAscope revealed that *C3*, *Cxcl12 and Fbln1* transcripts were localized to the media and the lesion cap within the atherosclerotic plaques. These findings suggest that E-cig aerosol exposure results in increased modulation of SMC resulting in an increased proportion of pro-inflammatory SMC subpopulation migrating into the atherosclerotic lesion, contributing to the inflammatory microenvironment within the plaque. To functionally validate the inflammatory SMC phenotype as suggested by our single-cell analysis, we performed a monocyte transwell migration assay using human coronary artery SMCs (HCASMCs) co-culture. E-cig aerosol-treated HCASMCs promoted increased monocyte migration, compared to E-cig or HCASMCs alone. This effect was attenuated by siRNA-mediated knockdown of C3, a marker of the pro-inflammatory FMC2 cluster, suggesting that this SMC cell type may contribute to E-cig-induced chemotactic signaling (**Figure S2F**).

### E-cig induces a global activation of chromatin accessibility

Our data indicates that E-cig aerosol exposure facilitates significant gene expression changes across multiple vascular cell types within the vessel wall with a notable effect in the vascular SMC. To further define the global epigenomic effects of E-cig aerosol exposure that may facilitate our observed transcriptomic and phenotypic changes, we performed nuclear isolation, 10X capture, and subsequent single cell ATAC sequencing (scATACseq) from the same single cell suspension as our scRNAseq analyses. Across a total of 5 captures (2 Filtered air, 3 E-cig; 1:1 mix of tdTomato-positive and tdTomato-negative cells), we performed peak calling and data aggregation (aggr) with CellRanger with downstream data quality evaluation with Signac and Seurat as we have previously done (see Methods)^21^. With a total of 66,890 nuclei that met our quality control, we performed UMAP dimensionality reduction and clustering, computed gene activity, and further tested the differential peak calling, gene activity, and transcription factor motif enrichment by exposure condition.

UMAP visualization grouped by exposure revealed distinct differences between Filtered air and E-cig (**Figure 3A**), and gene activity score of canonical genes allowed identification of SMC (*Myh11*), Fibroblast (*Pi16*), Macrophage (*Cd68*), and Endothelial (*Cdh5*) cell clusters (**Figure 3B**). Differential peak accessibility testing between exposure conditions revealed marked epigenomic changes where E-cig resulted in net increase in global chromatin accessibility in all cell types with the SMC and macrophage response being most pronounced (**Figure 3C, S3A**) (E-cig induced 3,598 differentially accessible chromatin peaks in SMCs, 3,807 in macrophage cells, 3,285 in fibroblasts, and 873 in endothelial cells (all P val adj <0.05, Log2 FC >0.25)). Visualization of top differentially accessible peaks highlighted activation of distinct enhancer elements (**Figures 3D-F**). Differentially accessible peaks mapped to genes such as *Zbtb2* (**Figure 3D**) with broad functions in transcriptional regulation, however additional top peaks mapped to key transcription factors involved in SMC phenotypic modulation such as *Egr1* and *Klf4* (**Figures 3E,F**). To further explore the global epigenomic effect of E-cig aerosol exposure on the vessel wall SMCs, we interrogated the peak to gene relationship and biological effect of the differentially accessible peaks of SMC-lineage cells with Genomic Regions Enrichment of Annotations Tool (GREAT).^25^ Top gene ontology molecular function included ‘unfolded protein binding’ and ‘mitogen-activated protein kinase binding’ with biological processes highlighting ‘regulation of intrinsic apoptotic signaling’, suggesting unfolded protein and cellular response to stress pathways (**Figures S3B,C**). Top gene ontology biological processes for macrophage cells include ‘immune response’, ‘positive regulation of immune system process’, ‘response to external biotic stimulus’ as well as ‘cell activation’ (**Figure S4A**). GO biological processes for fibroblasts and endothelial cells show some similarities including ‘regulation of gene expression/translation’, ‘regulation of cellular amide metabolic process’, and ‘mRNA catabolic process’ as shared processes within both cell types (**Figures S4B-C**).

**Figure 3.**
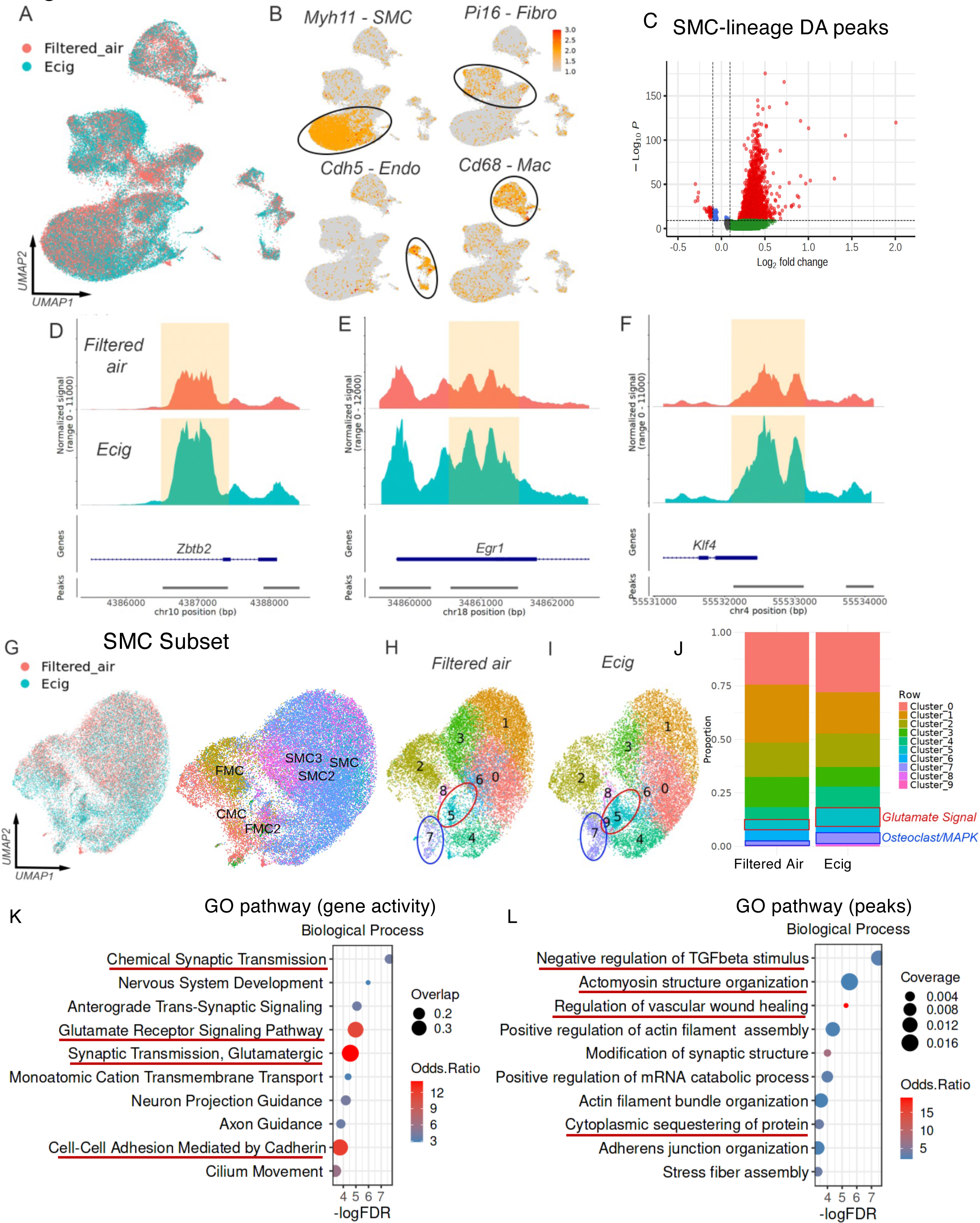
E-cig induces a global activation of chromatin accessibility. (A) UMAP of scATACseq data from atherosclerotic aortic root at 12 weeks of high fat diet with E-cig exposure in SMC lineage tracing mice (SMC-LnT: *Myh11^CreERT^*^2^, *ROSAtdTomato*, *ApoE^-/-^*) grouped by exposure type. (B) Feature plots of gene activity for canonical cell type markers *Myh11* – SMC, *Pi16* – Fibro, *Cdh5* – Endo, and *Cd68* – Mac. (C) Volcano plot of differentially accessible peaks by E-cig exposure within SMC subset. Coverage plots for Zbtb2 (D), Egr1 (E), and Klf4 (F) genomic regions split by exposure within SMC subset. In SMC subset, (G) UMAP visualization of scATACseq data grouped by exposure type (left) and integrated scRNAseq label transfer (right). UMAP of SMC subset with ATAC clusters in (H) filtered air and (I) E-cig exposed groups. (J) Stacked bar charts showing proportion of cells within SMC scATACseq clusters split by exposure. (K) GO Pathway analysis of top marker genes by gene activity for cluster ‘5’ in scATACseq SMC subset. (L) GO Pathway analysis of top marker genes by chromatin activity for cluster ‘5’ in scATACseq SMC subset.

### E-cig epigenetically activates glutamate neurotransmitter receptors and calcium signaling pathways in SMC

Unbiased clustering analysis of the scATAC-seq data revealed distinct cell type specific clusters including 5 SMC clusters (**Figure S5A)**. We performed a subsetted analysis of the SMC clusters with higher resolution to further understand the epigenomic changes within the SMC. With dimensionality reduction, UMAP data visualization, and clustering analysis, we identified 10 distinct clusters (**Figure S5A)**. Label transfer from scRNA-seq data identified the corresponding SMC cell types in the scATAC-seq clusters (**Figure 3G**). Consistent with the scRNAseq data, we found E-cig to induce a notable change in SMC cluster proportions (**Figures 3H-J**). Featureplot of *Myh11* chromatin activity representing contractile SMC phenotype (**Figure S5A**) identified contractile SMC clusters (clusters ‘1’, ‘0’, and ‘3’) as well as a modulated cluster (‘2’). Pathway analysis of the top genes for chromatin accessibility in cluster ‘2’ enriched for high ‘focal adhesion’ and ‘Adherens junction’ and ‘calcium signaling’ more consistent with an FMC/CMC phenotype (**Figure S5B**).

Distinct in the scATACseq data, however, we found additional clusters arose that appear biologically distinct from our previously described SMC/FMC/CMC phenotypic transition. Further evaluation identified E-cig to increase proportions of cluster ‘5’ and ‘7’ by 2.0-fold and 3.2-fold respectively (cluster ‘5’, FA 4.4%, E-cig 9.0%, *p<E-16*; cluster ‘7’, FA 1.7%, E-cig 5.4%, *p<E-16*) (**Figure 3J**). Notably, FindMarker analysis of gene activity revealed that cluster ‘5’ is marked by glutamate and neurological signaling pathways such as ‘Synaptic transmission’, ‘Glutamate receptor signaling pathway’ (**Figure 3K, S5C**). Key genes that drive these pathways include glutamate receptor genes (i.e. *Grin2a*/*2b/3a*, *Gria1*-*4*) and GABA receptors (i.e. *Gabra2/5, Gabrb2/3*). Additionally, analysis of the differentially accessible peaks in cluster ‘5’ using GREAT tools identified pathways relevant to atherosclerosis including regulation of TGF-β stimulus, vascular wound healing, and actomyosin structure organization (**Figure 3L**). Cluster ‘7’ is marked by ‘Osteoclast differentiation’, ‘MAPK signaling’, and ‘Fluid shear stress and atherosclerosis’ (**Figure S5D**). These data suggest that E-cig induces an epigenetic effect within the SMC that alters SMC phenotype and uniquely identifies glutamatergic and other neurological signaling pathways as potentially mediating this effect.

### E-cig activates transcription factor motif accessibility in vascular SMC

To understand how E-cig aerosol exposure may govern transcriptional gene programs by regulating transcription factor activation, we inferred transcription factor motif accessibility on single cell resolution with ChromVar^26^. Within the SMC subset, we identified that E-cig aerosol exposure activates TF motif accessibility for key regulatory TFs including GC rich family of motifs including *SP4* (MA0685.1), *KLF14* (MA0740.1) and *KLF16* (MA0741.1) (i.e. *SP4*, **Figures S6A-C**), as well as TFs with known function in SMC phenotypic transition including *KLF4* (MA0039.4), *EGR1* (MA0162.4), and *MEF2* family (MEF2A MA0052.4) (**Figures S6D-L**). To further understand the distinct relative TF motif accessibility for the newly identified ‘neurotransmitter’ cluster, Findmarker analysis identified this cluster to be enriched for interferon responsive TF motifs such as *IRF8* (MA0652.1), STAT signaling (*STAT5A*), ZBTB family of transcription factors (*ZBTB7A*, MA0750.2), as well as retinoid X receptors (*RXRB/G*, MA0855.1/MA0856.1) (**Figures S6M-O**).

### *Grin2a* is expressed by SMC population at the FMC and CMC junction

The observation of a distinct SMC cluster marked by glutamatergic signaling genes activated by E-cig suggested a potential relationship between activation of this pathway with formation of CMC in atherosclerosis. Further interrogation of the RNA expression of these glutamatergic NMDA and AMPA receptor subunit genes within the vascular SMC by scRNAseq revealed that the NMDA subunit *Grin2a* is increased in expression with Ecig exposure in the SMC population (**Figure 4A**) and is specifically expressed at the intersection of FMC to CMC transition (**Figure 4B**), suggesting a potential role in the phenotypic modulation of SMCs in response to E-cig aerosol exposure. Coverage plot of representative glutamate receptor gene *Grin2a* suggests E-cig to activate regulatory enhancers of *Grin2a* (**Figure S7A**). Furthermore, Spearman correlation analysis revealed that *Grin2a* expression is significantly correlated with FMC and CMC markers such as *Lum*, *Spp1*, and *Runx1*, while exhibiting negative correlations with SMC-specific markers including *Myh11*, *Tagln*, *Cnn1*, and *Acta2* (**Figure 4C**).

**Figure 4.**
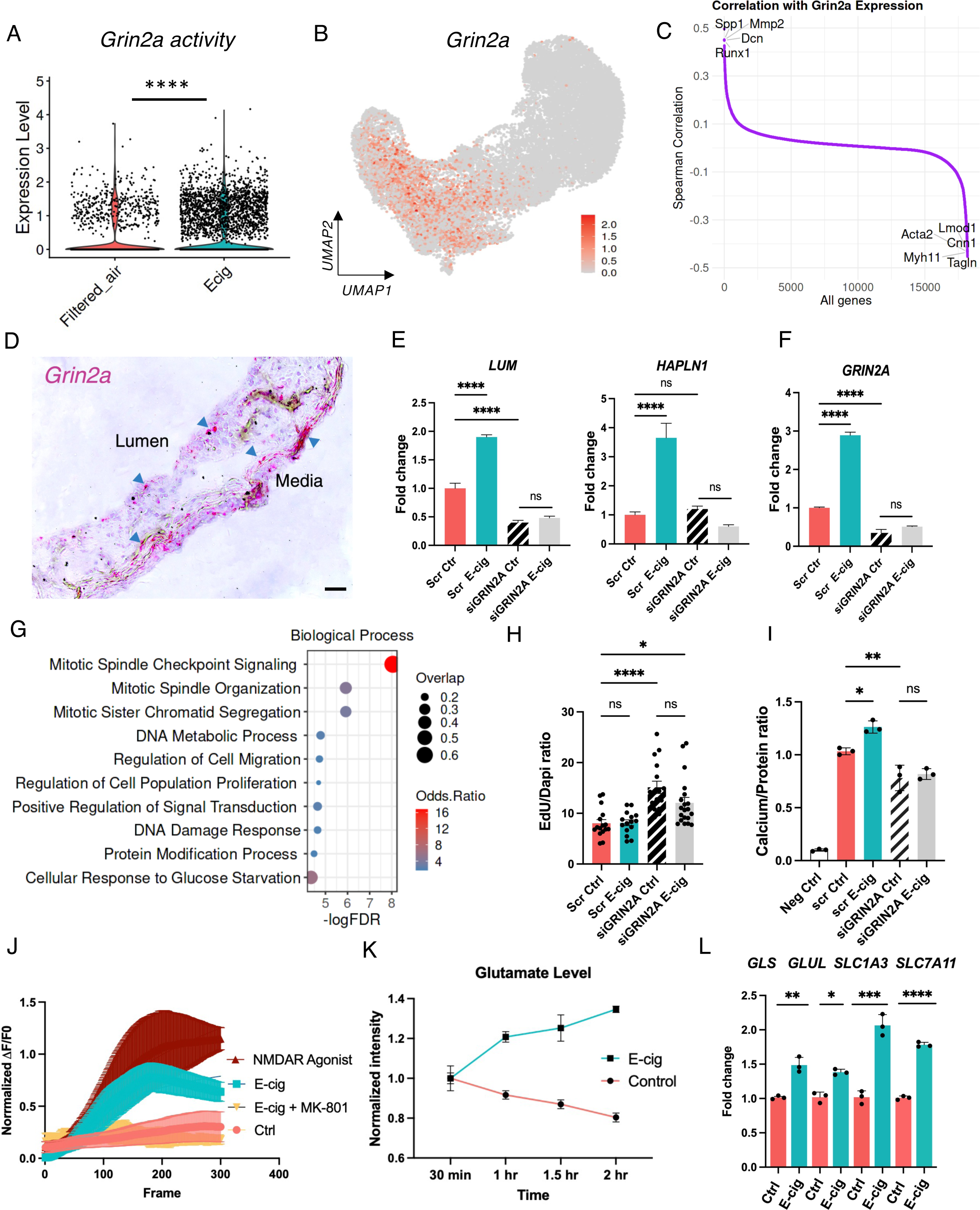
***GRIN2A* expression defines a phenotypically modulated SMC subset at the FMC–CMC junction and links E-cig exposure to altered glutamate metabolism and SMC remodeling.** (A) Violin plot depicting the distribution of *Grin2a* expression in the SMC subset from Filtered air–exposed and E-cig–exposed groups. E-cig exposure was associated with significantly higher expression compared with Filtered air control, with p-value obtained using two-tailed T-test, *P* < 0.0001. (B) Expression *Grin2a* across FMC and CMC junction by scRNA-seq. (C) Spearman correlation analysis of *GRIN2A* with every other gene in cells across the ‘SMC’, ‘FMC’, ‘CMC’ clusters. Selected examples of *GRIN2A*-regulated genes are labeled. (D) RNAscope in situ hybridization of *GRIN2A* in the aortic root of E-cig exposed mice. The image is representative of three experiments, and the scale bar represents 50 μm. Blue arrows highlight cells at the lesion and the media. (E) Expression of FMC (*LUM)* and CMC (*HAPLN1*) markers were evaluated by quantitative RT-PCR with *GRIN2A* knockdown in HCASMCs. Data were expressed as mean +/- SEM. With p-value obtained using one-way ANOVA with Tukey’s multiple comparisons post-hoc test, For *LUM* (scrambled Ctrl vs scrambled E-cig P<0.0001, scrambled Ctrl vs si*GRIN2A* Ctrl P<0.0001, siG*RIN2A* Ctrl vs si*GRIN2A* E-cig P=0.2252) For *HAPLN1* (scrambled Ctrl vs scrambled E-cig P<0.0001, scrambled Ctrl vs si*GRIN2A* Ctrl P=0.69, si*GRIN2A* Ctrl vs si*GRIN2A* E-cig P=0.0083). Dots represent three technical replicates from three biologically independent samples. (F) Expression of *GRIN2A* was evaluated in HCASMCs by quantitative RT-PCR with *GRIN2A* knockdown in HCASMC. Data were expressed as mean +/- SEM. With p-value obtained using one-way ANOVA with Tukey’s multiple comparisons post-hoc test, (scrambled Ctrl vs scrambled E-cig P<0.0001, scrambled Ctrl vs si*GRIN2A* Ctrl P<0.0001, si*GRIN2A* Ctrl vs si*GRIN2A* E-cig P=0.052). Dots represent three technical replicates from three biologically independent samples. (G) GO pathway enrichment analysis from differentially expressed genes in si*GRIN2A* Ctrl versus scrambled Ctrl samples. (H) EdU incorporation assay used to measure proliferation of HCASMCs with *GRIN2A* knockdown. Data were expressed as mean +/- SEM. With p-value obtained using one-way ANOVA with Dunnett’s multiple comparisons post-hoc test, (scrambled Ctrl vs scrambled E-cig P=0.99, scrambled Ctrl vs si*GRIN2A* Ctrl P<0.0001, scrambled Ctrl vs si*GRIN2A* E-cig P=0.035, si*GRIN2A* Ctrl vs si*GRIN2A* E-cig P=0.056). (I) Rate of calcification with *GRIN2A* knockdown in HCASMCs grown in calcification media. Data were expressed as mean +/- SEM. With p-value obtained using one-way ANOVA with Tukey’s multiple comparisons post-hoc test, (scrambled Ctrl vs scrambled E-cig P=0.011, scrambled Ctrl vs si*GRIN2A* Ctrl P=0.0059, si*GRIN2A* Ctrl vs si*GRIN2A* E-cig P=0.96). Dots represent three technical replicates from three biologically independent samples. Neg Ctrl = cells cultured in the absence of calcification media. (J) Real-time calcium influx in HCASMCs measured using the FLUO-4 AM probe following treatment with E-cig extract, MK-801 (NMDAR antagonist) plus E-cig extract, or NMDA plus D-serine (NMDAR agonists). Calcium dynamics were recorded immediately after treatment for 300 seconds using an ECHO Revolve microscope. Data are expressed as ΔF/F0 where F0 is baseline fluorescence intensity of the first 10 frames and ΔF is the change in mean fluorescence intensity. (K) Extracellular glutamate was quantified into the supernatant of HCASMCs following E-cig treatment. Data were normalized relative to Ctrl and expressed as mean +/- SEM. With p-value obtained using two-way ANOVA with Sidak’s multiple comparisons post-hoc test, (60min P<0.0001, 90min P<0.0001, 120 min P<0.0001). (L) Expression of glutamate-metabolizing enzymes (*GLS, GLUL*) and amino acid transporters (*SLC1A3*, *SLC7A11*) were evaluated in HCASMCs by quantitative RT-PCR. Data were normalized relative to Ctrl and expressed as mean +/- SEM. With p-value obtained using two-tailed T-test, (*GLS* P=0.0021; *GLUL* P=0.048; *SLC1A3* P=0.0006; *SLC7A11* P<0.0001). Dots represent three technical replicates from three biologically independent samples.

**Figure 5.**
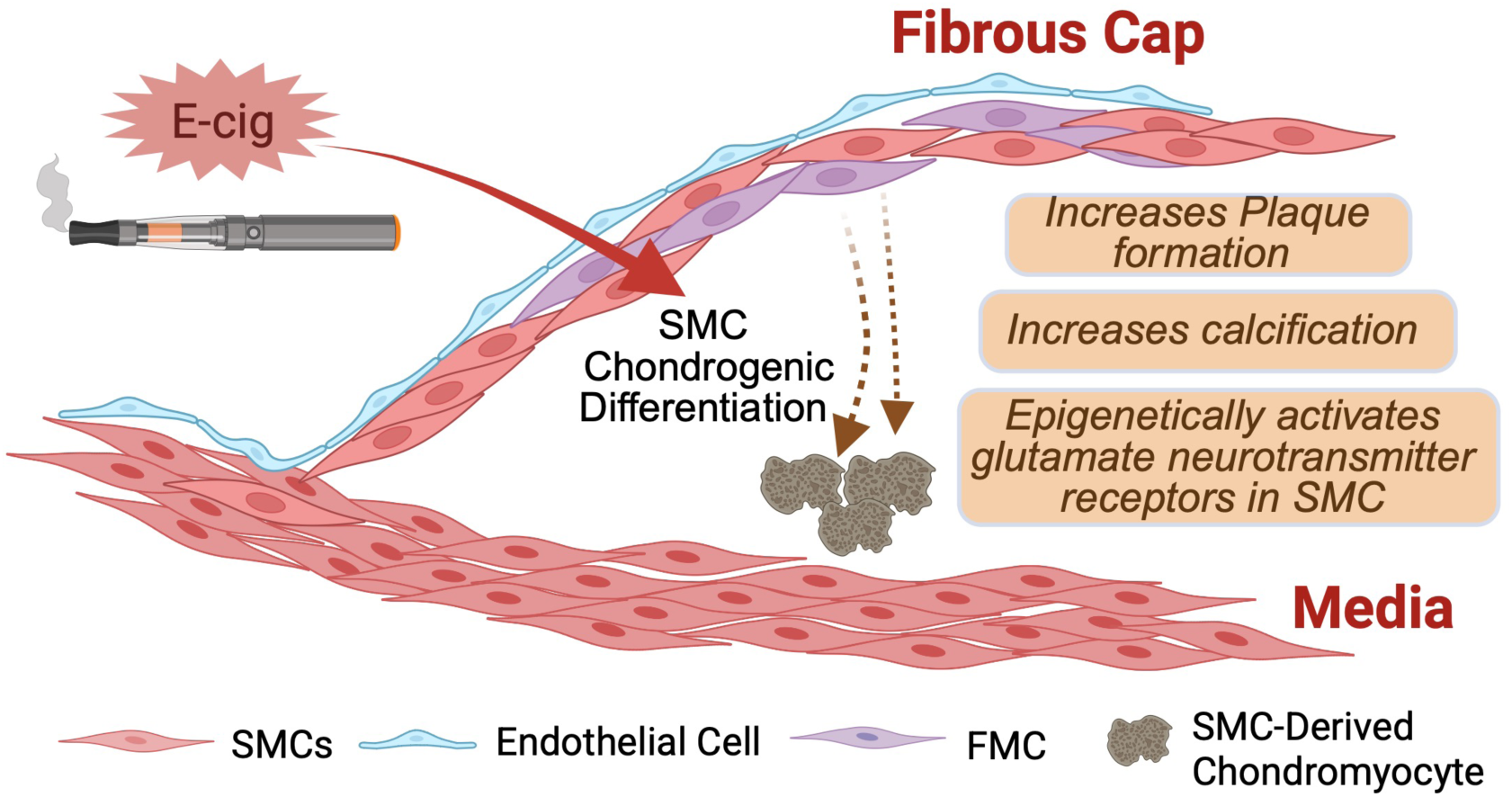
Graphical Abstract. E-cig exposure drives smooth muscle cell modulation toward a chondrogenic state, promoting atherosclerosis progression and calcified plaque formation. These pathogenic changes are mediated by coordinated transcriptional reprogramming and epigenetic remodeling, including activation of NMDA–glutamate receptor pathways.

To further understand where the *Grin2a* expressing cells reside spatially within atherosclerotic plaque, we performed RNA scope in-situ hybridization that revealed *Grin2a* expression to be localized to the media and the lesion cap within the atherosclerotic plaques (**Figure 4D**). Together, these data suggest that *Grin2a* expression is augmented by E-cig aerosol exposure and may play a role in mediating SMC phenotypic transition from FMC to CMC observed *in vivo*.

### E-cig aerosol exposure drives phenotypic modulation and *GRIN2A* expression in HCASMCs

Building on our *in vivo* findings implicating a link between E-cig aerosol exposure, *GRIN2A*, and smooth muscle cell (SMC) phenotypic modulation, we investigated the direct effects of E-cig and *GRIN2A* signaling in HCASMCs *in vitro*. E-cig extract was generated from a commercially available JUUL device, and the concentration used was selected based on preliminary cytotoxicity and functional assays, which confirmed it to be non-toxic yet biologically active (**Figure S7B**; see Methods for details).

To assess how E-cig aerosol exposure alters SMC phenotype, we treated cultured HCASMCs with E-cig extract and measured expression of established markers of SMC dedifferentiation, including *LUM* and *HAPLN1*. E-cig treatment significantly upregulated both markers, indicating a shift toward a dedifferentiated SMC phenotype (**Figure 4E**).

Furthermore, we found that e-cig extract treatment significantly increased *GRIN2A* expression in HCASMC (**Figure 4F**), identifying this NMDAR subunit as a potential molecular target of E-cig aerosol exposure. To determine whether *GRIN2A* plays a functional role in mediating SMC phenotypic modulation, we modified *GRIN2A* expression in HCASMCs using siRNA knockdown (**Figures 4E-F**). Notably, knockdown of *GRIN2A* resulted in complete inhibition of the E-cig induced activation of SMC dedifferentiation markers, including *LUM* and *HAPLN1* (**Figure 4E**). Conversely, *GRIN2A* overexpression amplified the effects of E-cig treatment on SMC dedifferentiation markers (**Figure S7C**). These findings identify *GRIN2A* as a critical mediator of E-cig-induced SMC phenotypic modulation *in vitro* and position NMDAR signaling as a novel mechanistic pathway through which E-cig promote vascular remodeling.

### *GRIN2A* negatively regulates SMC proliferation and promotes chondrogenic gene programs and calcification *in vitro*

To explore the transcriptional pathways regulated by *GRIN2A* in SMC, we performed siRNA KD in HCASMC followed by bulk RNA sequencing. At an adjusted *P* value <0.05 and fold change >1.3, *GRIN2A* KD resulted led to widespread transcriptional changes, with 811 upregulated and 569 downregulated genes (**Figure S7D, S7E; Table S2**). GO enrichment analysis found *GRIN2A* KD to modify biological processes including cell cycle regulation, cell migration, and proliferation, protein modification response, and response to glucose starvation (**Figure 4G**).

To functionally validate the impact of *GRIN2A* on cell proliferation, we performed a 5-ethynyl-2′-deoxyuridine (EdU) proliferation assay (**Figure 4H**). Consistent with our transcriptomic findings, *GRIN2A* knockdown significantly increased HCASMC proliferation, independent of E-cig aerosol exposure. These results suggest that *GRIN2A* suppresses cell cycle progression, potentially promoting the transition from a proliferative SMC state to a more calcified phenotype.

To assess the role of *GRIN2A* activation in vascular calcification, we conducted *in vitro* calcification assays using HCASMC. Cells were cultured in calcification media for 6 days, after which *GRIN2A* knockdown significantly decreased the calcification phenotype **(Figure 4I),** while *GRIN2A* overexpression enhanced it (**Figure S7F**). These findings further support the role of *GRIN2A* as a key regulator of chondrogenic differentiation in SMCs.

### E-cig results in significant alteration in the glutamate metabolism and increases *Ca2+ flux*

Given the functional role of *GRIN2A* in HCASMC, we next examined whether NMDA receptors (NMDAR) are functional in SMC and if E-cig aerosol exposure can modulate NMDAR activity. NMDAR activation results in intracellular calcium flux and requires the simultaneous binding of two ligands: a primary agonist such as glutamate (or NMDA, a synthetic glutamate analog), and a co-agonist such as D-serine^27^. As a functional readout of NMDAR activation, we assessed intracellular calcium (Ca²⁺) flux using live-cell imaging. To avoid interference from glutamate present in standard culture media, cells were stimulated in a Mg²⁺-free HEPES-buffered saline solution devoid of glutamate. This solution also facilitates NMDAR activation by relieving voltage-dependent Mg²⁺ blockade of the receptor pore under resting membrane potential. Stimulation with NMDA and D-serine induced a robust increase in intracellular Ca²⁺ levels in HCASMCs, confirming the presence of functional NMDAR in HCASMCs (**Figure 4J**). E-cig treatment of HCASMC also resulted in significantly increased intracellular Ca²⁺ levels in HCASMCs, which was markedly attenuated by pre-treatment with MK-801, a non-competitive antagonist of NMDAR, supporting the functional activation of NMDAR by E-cig in HCASMC.

Next, we investigated whether E-cig aerosol exposure alters glutamate balance in SMCs, which could potentially lead to change in NMDAR activity. Treatment of HCASMC with E-cig extract for 2 hours led to a significant increase in extracellular glutamate levels in the culture supernatant, suggesting enhanced glutamate release or impaired reuptake mechanisms (**Figure 4K**). To assess whether these changes were associated with altered glutamate metabolism, we examined the expression of enzymes and transporters involved in the glutamate-glutamine cycle. E-cig treatment significantly upregulated glutaminase (*GLS1*), which catalyzes the conversion of glutamine to glutamate, and glutamate-ammonia ligase (*GLUL*), which synthesizes glutamine from glutamate. We also observed increased expression of *SLC1A3*, the gene encoding high-affinity sodium-dependent glutamate transporter EAAT1, which mediates glutamate reuptake and *SLC7A11* which encodes the cystine-glutamate antiporter xCT **(Figure 4L)**. These findings suggest that E-cig aerosol exposure perturbs glutamate homeostasis, potentially triggering a compensatory metabolic response aimed at rebalancing glutamate-glutamine flux.

To assess *GRIN2A* expression in human disease, RNAscope in situ hybridization was performed in coronary artery sections from heart transplant recipients with atherosclerosis. *GRIN2A* expression was detected in the lesion, with the strongest signal localized to the base of the intimal layer **(Figure S7G)**. In parallel, to define the spatial distribution of glutaminase expression, we performed RNAscope for *GLS1* in the same disease context **(Figure S7G).** *GLS1* expression was also enriched within the lesion at the basal intimal layer, consistent with increased glutaminase activity in human atherosclerotic plaques. Collectively, these findings indicate that human atherosclerotic lesions exhibit capacity for enhanced glutamate production alongside *GRIN2A* expression, and support a model in which environmental exposures may amplify engagement of NMDAR-linked signaling programs in vascular smooth muscle cells.

To further investigate if this pathway is mediated by nicotine, a major component of E-cig liquids, HCASMCs were treated with nicotine at a range of physiologically relevant concentrations. We measured both *GRIN2A* expression and intracellular calcium flux **(Figure S7H and S7I)**, however, nicotine did not induce *GRIN2A* expression, nor did it increase intracellular calcium flux, suggesting that nicotine alone is not sufficient in activating the glutamatergic signaling pathway.

## Discussion

E-cigarette (E-cig) use is rapidly increasing worldwide, yet its long-term cardiovascular health consequences remain incompletely understood. While inhaled toxicants, including cigarette smoke and fine particulate matter, are well-documented to affect vascular function and disease risk, the mechanisms by which E-cig aerosols influence vascular homeostasis and epigenomic regulation are less well defined. Given that E-cig aerosols deliver a complex mixture of chemicals that rapidly enter systemic circulation, it is plausible they interact directly with the vasculature to elicit pathogenic effects. Here, we expand the limited mechanistic literature on E-cig-induced vascular toxicity^28–32^, presenting the first study to profile the global transcriptomic and epigenomic landscape of the aortic wall at single-cell resolution following chronic E-cig aerosol exposure. Using genetic lineage tracing, we further dissected the specific contribution of vascular smooth muscle cells (SMCs) to this pathophysiological response.

Our findings align with previous studies showing that E-cig-aerosol exposure exacerbates atherosclerosis^28,31^. However, our data uniquely demonstrate that SMCs are the most responsive cell type at both the transcriptional and epigenetic levels. Single-cell RNA sequencing revealed marked transcriptional remodeling in SMCs, while endothelial cells, fibroblasts, and macrophages exhibited more modest changes. This underscores the importance of SMC phenotypic plasticity as a key determinant of environmental responsiveness in atherosclerotic disease^33,34^.

A key novel finding of our study is that chronic E-cig aerosol exposure accelerates atherosclerosis and promotes SMC phenotypic modulation toward a chondrogenic, calcifying state. This shift is characterized by downregulation of canonical contractile markers and upregulation of ossification-related pathways. These changes were accompanied by increased alkaline phosphatase activity *in vivo*, consistent with vascular calcification. Importantly, these effects occurred independently of systemic lipid changes, supporting a direct vascular specific mechanism of injury.

In addition to driving SMC chondrogenic differentiation, E-cig aerosol exposure induced a pro-inflammatory SMC phenotype. We observed increased macrophage recruitment in the lesion despite modest transcriptional responses in macrophages themselves. These results are in line with Caruana et al. who reported no significant changes in systemic immune cell profiles following E-cig aerosol exposure^30^. In our single-cell analysis, we identified a cluster of modulated SMCs (FMC2) that were marked by increased expression of genes including *Cxcl12*, *C3*, and *Fbln1* which have been known to increase the inflammatory milieu and chemotactic recruitment of inflammatory cells^35–38^. Further validation with *in vitro* transwell chemotactic assays suggests that this inflammatory SMC population may contribute to the increased macrophage recruitment. Further work will be required to assess their causal role in E-cig induced atherosclerosis.

We next examined how environmental exposures such as E-cig aerosol might epigenetically reprogram vascular cell states. Environmental toxicants are increasingly recognized to influence disease risk by reshaping cellular phenotype through alterations in global epigenomic signatures and transcriptional regulatory networks^39,40^. These regulatory networks, encompassing chromatin accessibility, transcription factor (TF) activity, and gene expression, underpin key mechanisms of vascular disease susceptibility^41^, and accumulating evidence suggests that environmental exposures can dynamically reconfigure these networks^40^. In our hyperlipidemia model of atherosclerosis, we performed single-cell chromatin accessibility profiling (scATAC-seq) of aortic tissue following E-cig aerosol exposure and uncovered substantial global remodeling of the epigenome. E-cig aerosol exposure induced a net increase in chromatin accessibility, particularly within SMCs. We observed enrichment of TF motifs associated with *Klf4*, *Egr1*, and *Mef2*, regulators known to control SMC plasticity and phenotypic switching. These data indicate that E-cig aerosol exposure activates transcriptional circuits that govern SMC state transitions and lesion remodeling.

One particularly intriguing finding of this study was the epigenetic activation of a glutamatergic signaling axis within SMCs. We observed increased accessibility and expression of *Grin2a* and *Grin2b*, which encode subunits of the NMDA (*N*-methyl-D-aspartate) receptor (NMDAR). NMDARs are ionotropic glutamate receptors that play a key role in synaptic plasticity, learning, and memory, primarily through their function as Ca²⁺-permeable channels in neurons^42^. While traditionally associated with the central nervous system, NMDAR expression has also been detected in peripheral tissues, including vascular smooth muscle cells and endothelial cells^43–45^. However, the functional role of NMDARs in vascular biology remains poorly defined. In our study, these epigenetic changes were concentrated in modulated SMCs transitioning toward a chondromyocyte (CMC) phenotype. Although *Grin2a* is not uniquely induced by E-cig aerosols, our data suggest increased glutamatergic signaling, potentially through amplification of endogenous glutamate pathways.

To mechanistically explore this glutamatergic signaling axis, we used *in vitro* models of HCASMCs. Functional experiments confirmed that *GRIN2A* is induced by E-cig aerosol exposure and is required for the E-cig-induced SMC phenotypic modulation. *GRIN2A* promoted chondrogenic gene expression, suppressed proliferation, and enhanced calcification. Parallel experiments showed that *GRIN2A* overexpression promoted SMC dedifferentiation and calcification, while knockdown reversed these phenotypes, confirming its role as a key regulator of this remodeling process. Furthermore, E-cig extract increased extracellular glutamate levels, and triggered calcium influx via activation of NMDARs. These findings support a model in which E-cig aerosol exposure induces dysregulation of the glutamate-glutamine cycle, leading to glutamate accumulation in the extracellular space and activation of NMDAR-dependent calcium signaling in vascular SMCs. Importantly, this calcium influx was abrogated by pharmacologic NMDAR inhibition and potentiated by co-agonists, confirming functional receptor activity in SMCs.

Although nicotine is a prominent and well-characterized component of E-cig aerosol, treatment of HCASMCs with nicotine alone, across a broad range of physiologically relevant concentrations, was not sufficient to induce significant *GRIN2A* expression or calcium influx. This suggests that other aerosol constituents, such as aldehydes, reactive oxygen species, flavoring chemicals, or heavy metals, may act independently or synergistically to initiate glutamate signaling in the vascular wall. Given the extensive chemical diversity and variability among E-cig products, further studies will be essential to dissect the contribution of individual toxicants to this pathway.

Our findings position *GRIN2A*-mediated NMDAR signaling as a contributory regulator of SMC phenotypic modulation and vascular remodeling. We propose that sustained calcium influx through NMDARs may activate downstream transcriptional programs governed by calcium-sensitive factors such as MEF2, a known regulator of vascular SMC plasticity^46,47^. Furthermore, chronic calcium elevation may contribute to mitochondrial dysfunction, oxidative stress, or endoplasmic reticulum (ER) stress, all of which are implicated in osteochondrogenic differentiation and vascular calcification^48–50^. These findings provide a mechanistic framework linking environmental toxicant exposure to vascular disease through glutamatergic signaling and underscore the vulnerability of vascular SMCs to non-nicotine aerosol constituents.

While NMDARs are well-characterized in neurons^51,52^, their roles in vascular SMCs are only beginning to be understood. Previous studies have identified endogenous NMDAR expression in pulmonary artery SMCs, where glutamate-NMDAR signaling has been shown to promote proliferation and contribute to the development of pulmonary arterial hypertension (PAH)^44,45^. However, the function of NMDARs in systemic vascular SMCs and their response to environmental stimuli remain poorly defined. Our study builds on this limited literature by demonstrating that human coronary artery SMCs endogenously express functional NMDARs and that this pathway is epigenetically activated in response to E-cig aerosol exposure. Notably, in contrast to the proliferative role described in pulmonary vessels, we observed that NMDAR activation in SMCs suppresses proliferation and instead promotes phenotypic modulation toward a chondrogenic, calcifying state. These findings suggest that NMDAR signaling exerts vascular bed- and context-specific effects, potentially governed by differences in the local environment, availability of co-agonists, or engagement of distinct downstream transcriptional programs. Our work expands the understanding of glutamatergic signaling in the vasculature and implicates this pathway as a novel mediator of environmentally induced vascular remodeling.

Several limitations of our study should be acknowledged. First, we employed an SMC-lineage tracing model, which specifically targets analysis of smooth muscle cell (SMC) phenotypic modulation during atherosclerosis. While our single-cell transcriptomic data permitted comparisons across multiple vascular cell types, the ability to comprehensively assess contributions from non-SMC populations within the vessel wall remains limited. Additionally, our animal studies were restricted to male mice due to the presence of the *Myh11-Cre* transgene on the Y chromosome. Given prior evidence of sex-dependent effects of E-cig aerosol exposure on atherosclerosis, future studies will be necessary to explore potential sex-based differences in SMC phenotypic modulation and glutamatergic signaling in response to E-cig aerosol exposure. Lastly, while our data establish *GRIN2A* as a key regulator of SMC phenotypic modulation, the upstream regulatory pathways that govern *GRIN2A* expression remain unclear. Future studies should investigate the epigenetic drivers and transcription factors involved in *GRIN2A* activation.

In conclusion, this study contributes to the growing body of evidence supporting E-cig aerosol exposure as a potent inducer of vascular remodeling and atherosclerosis and identify a previously unrecognized role for glutamate metabolism and NMDAR signaling in mediating E-cig-induced vascular remodeling. Through integration of single-cell and functional analyses, we show that this pathway promotes phenotypic modulation of SMCs toward a calcifying, chondrogenic fate. These findings not only expand our understanding of how environmental toxicants impact vascular biology but also reveal a previously unappreciated role for neurotransmitter signaling pathways in cardiovascular disease, supported by evidence of glutamate metabolic activity and *GRIN2A* expression in human atherosclerotic plaques.

### Source of Funding

This work is made possible by the generosity of our funders. National Institutes of Health grants K08HL153798 (P.C.), K08HL153798-S1(HuBMAP)(P.C.), R01HL151535 (J.B.K), L30HL159413 (C.W.), K08HL167699 (C.W.), R01 DC020176 (E.E.D); and American Heart Association grants 24POST1193717 (I.D.), 24POST1189844 (G.Q.), 23SCISA1144703 (P.C.), 24SCEFIA1248386 (P.C.), 20CDA35310303 (P.C.), 24POST1187860 (J.P.M.), 23CDA1042900 (C.W.); the Tobacco-Related Disease Research Program T32IR5352 (J.B.K), T32IR5240 (J.B.K).

## Acknowledgements

We gratefully acknowledge the Stanford Department of Cardiothoracic Surgery Human Biorepository Tissue Bank, especially Claire DaValle, for their assistance with the collection of human tissue.

