## Supplemental Figures for "Chronic Electronic Cigarette Exposure Promotes Atherosclerosis and Chondrogenic Modulation of Smooth Muscle Cells"

### Supplemental Figure 1

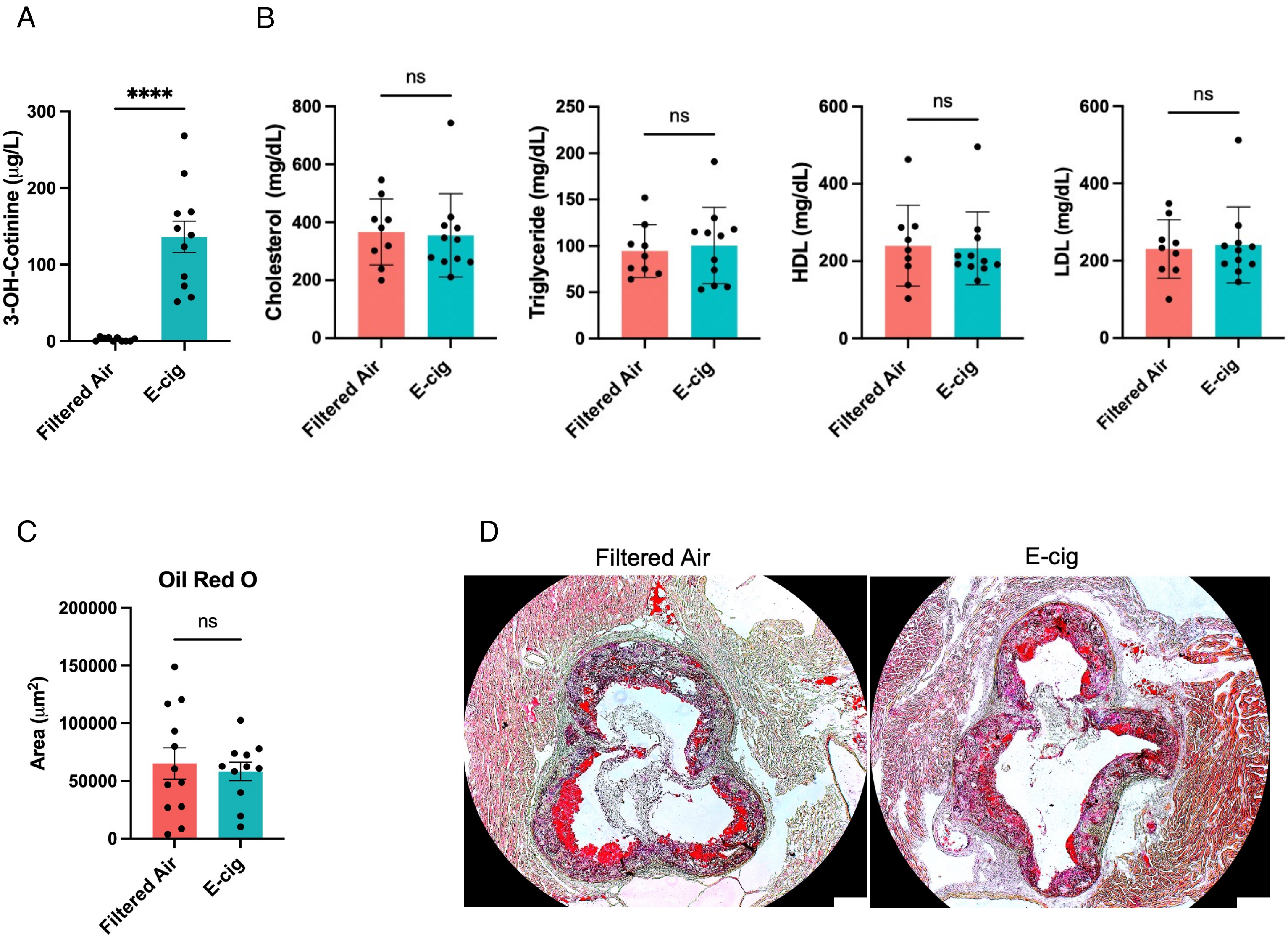

### Supplemental Figure 2

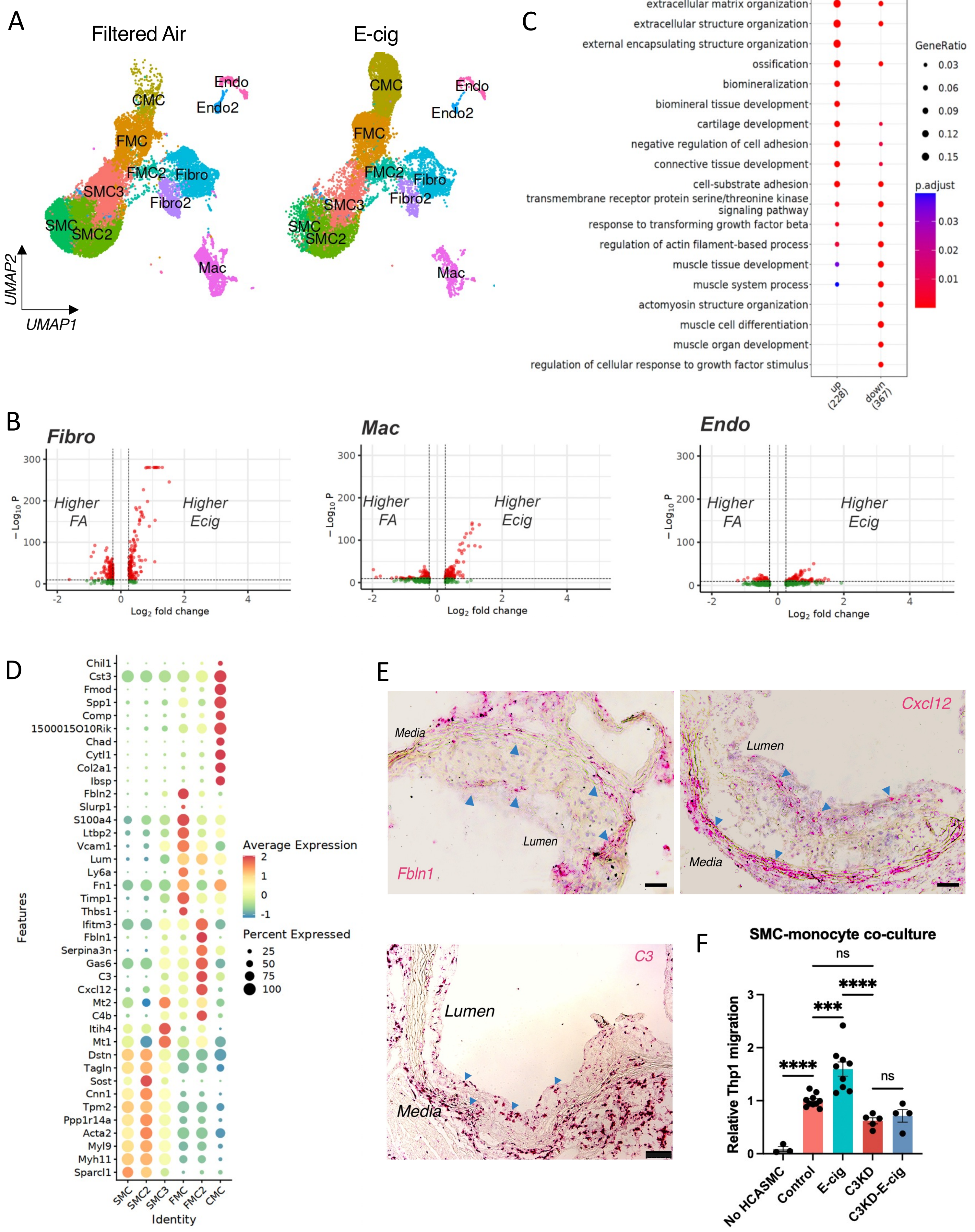

Supplemental Figure 3

A

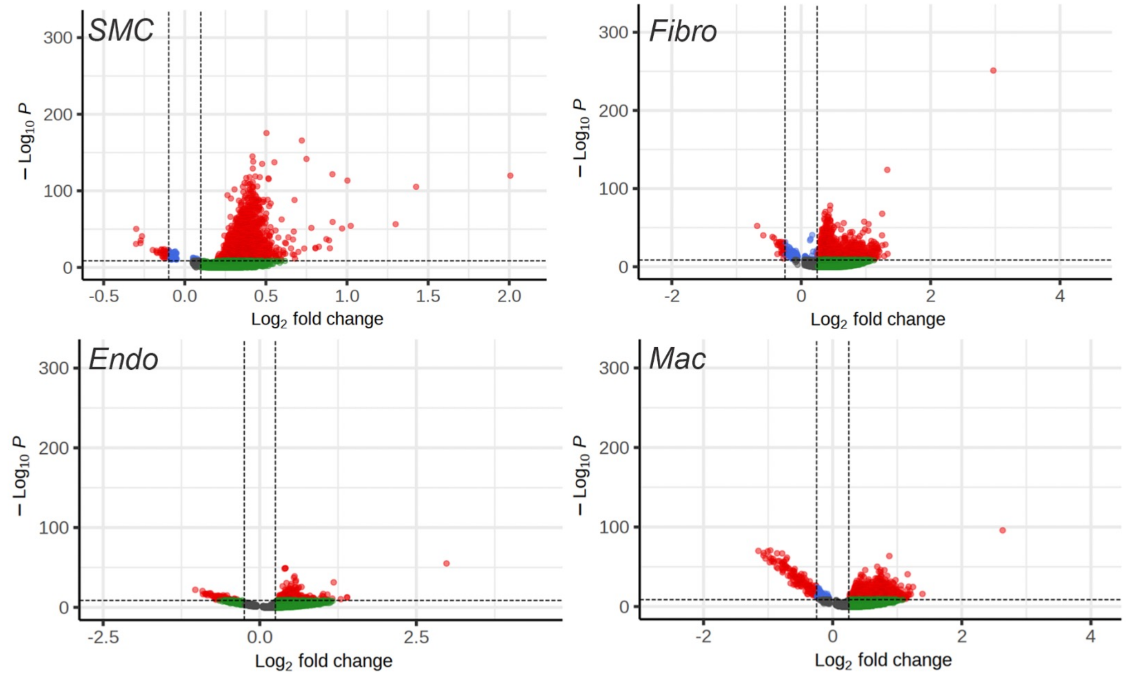


B

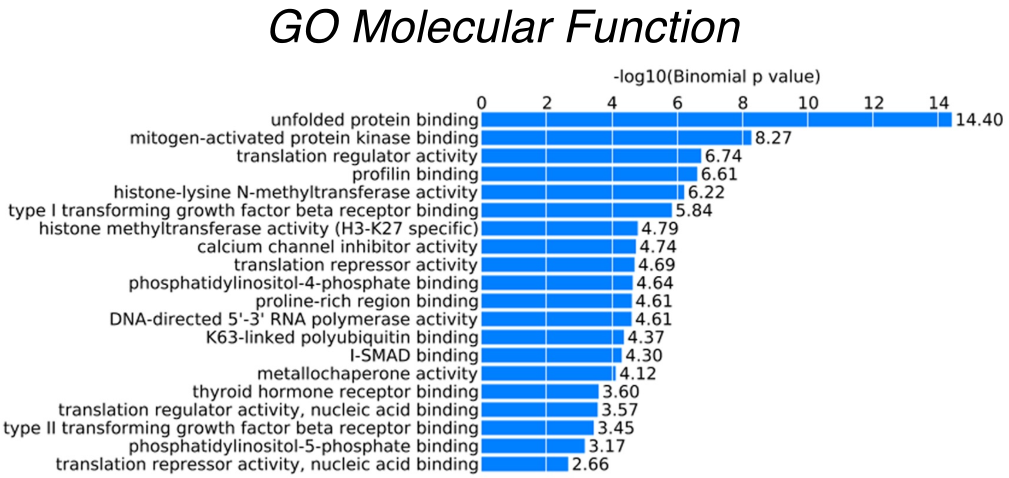


C

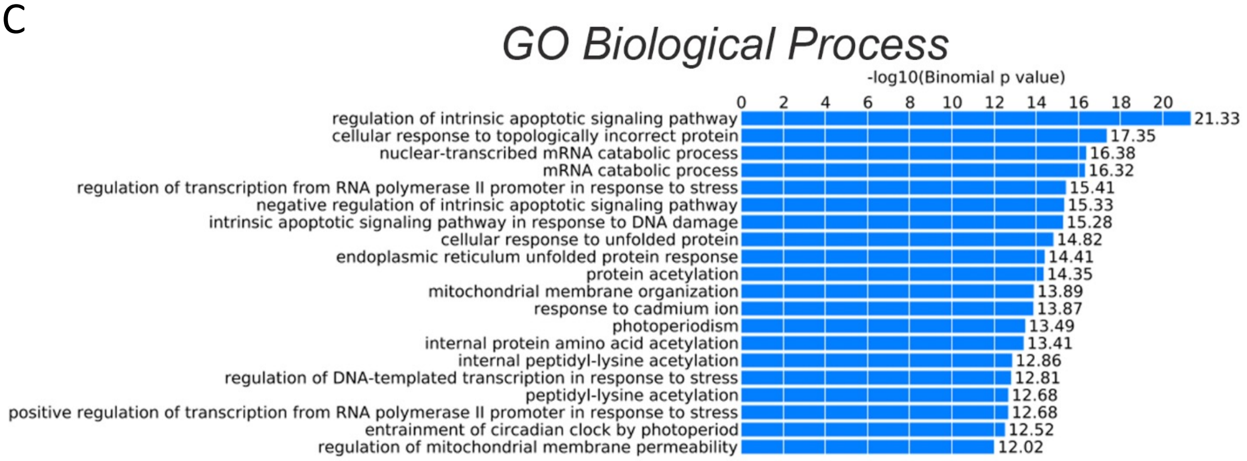

Supplemental Figure 4

A

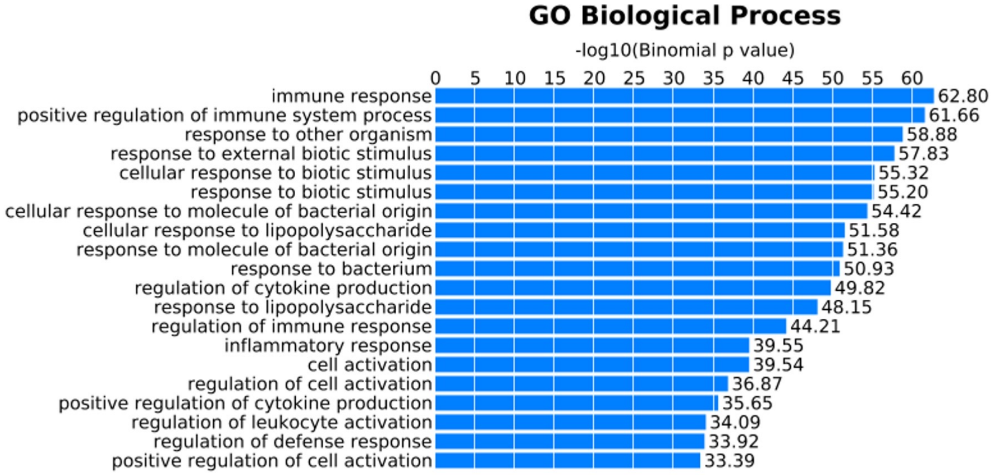


B

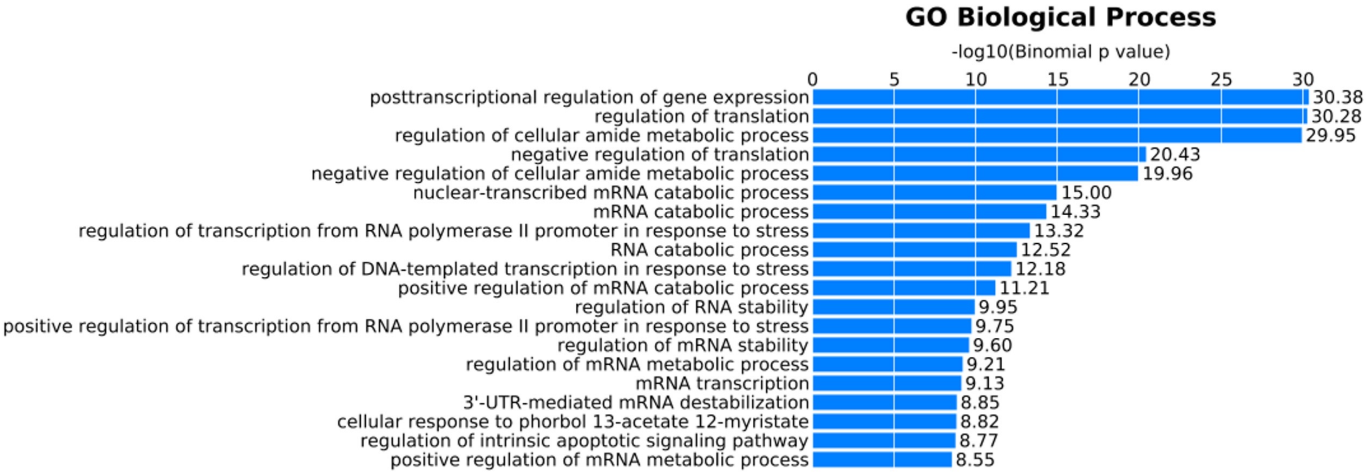


C

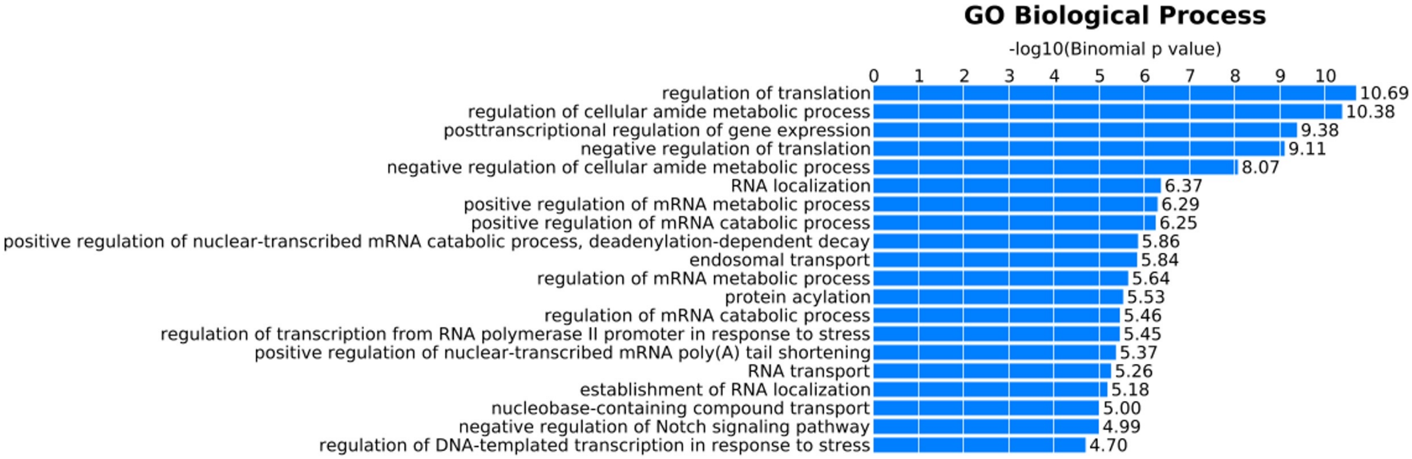

### Supplemental Figure 5

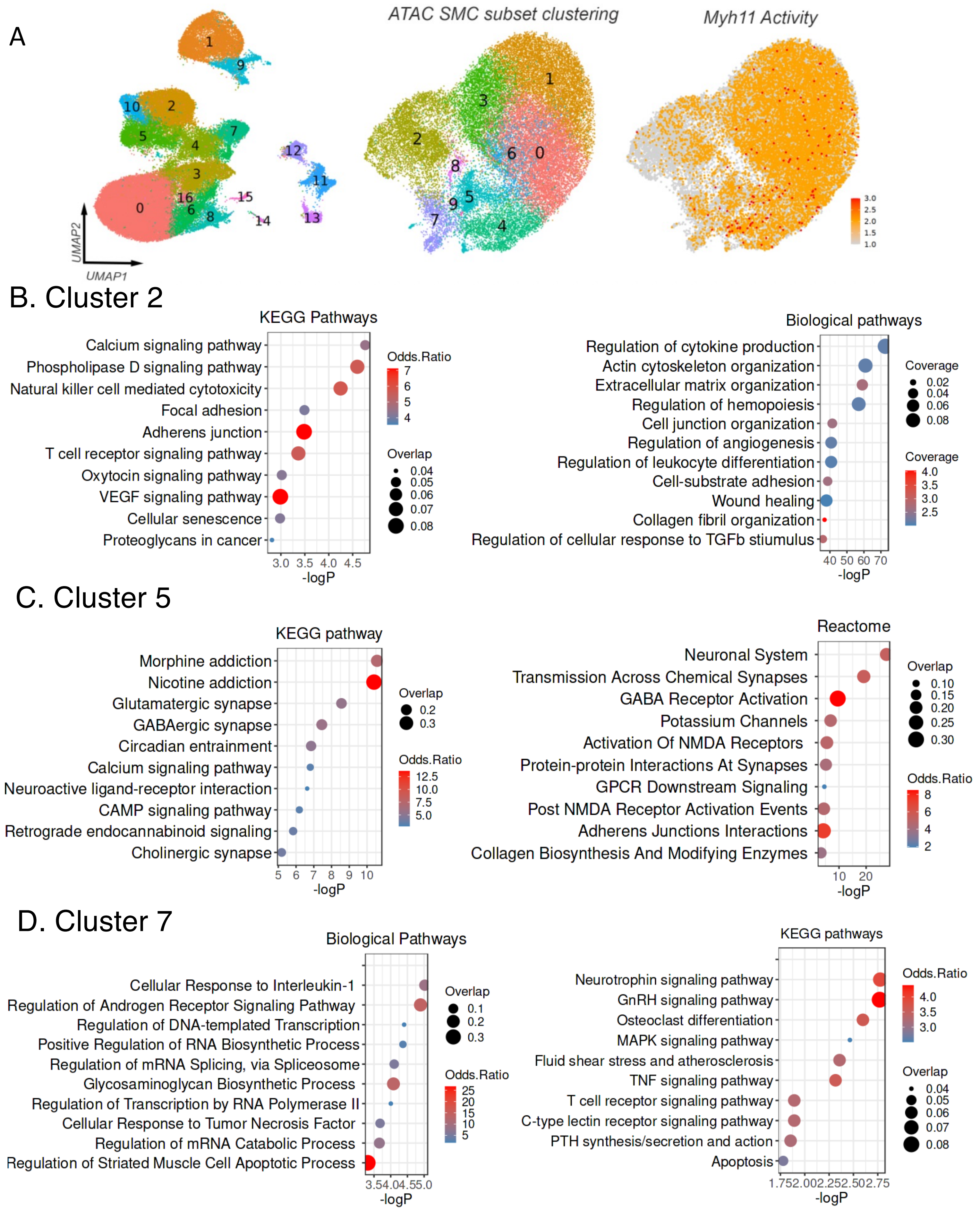

### Supplemental Figure 6

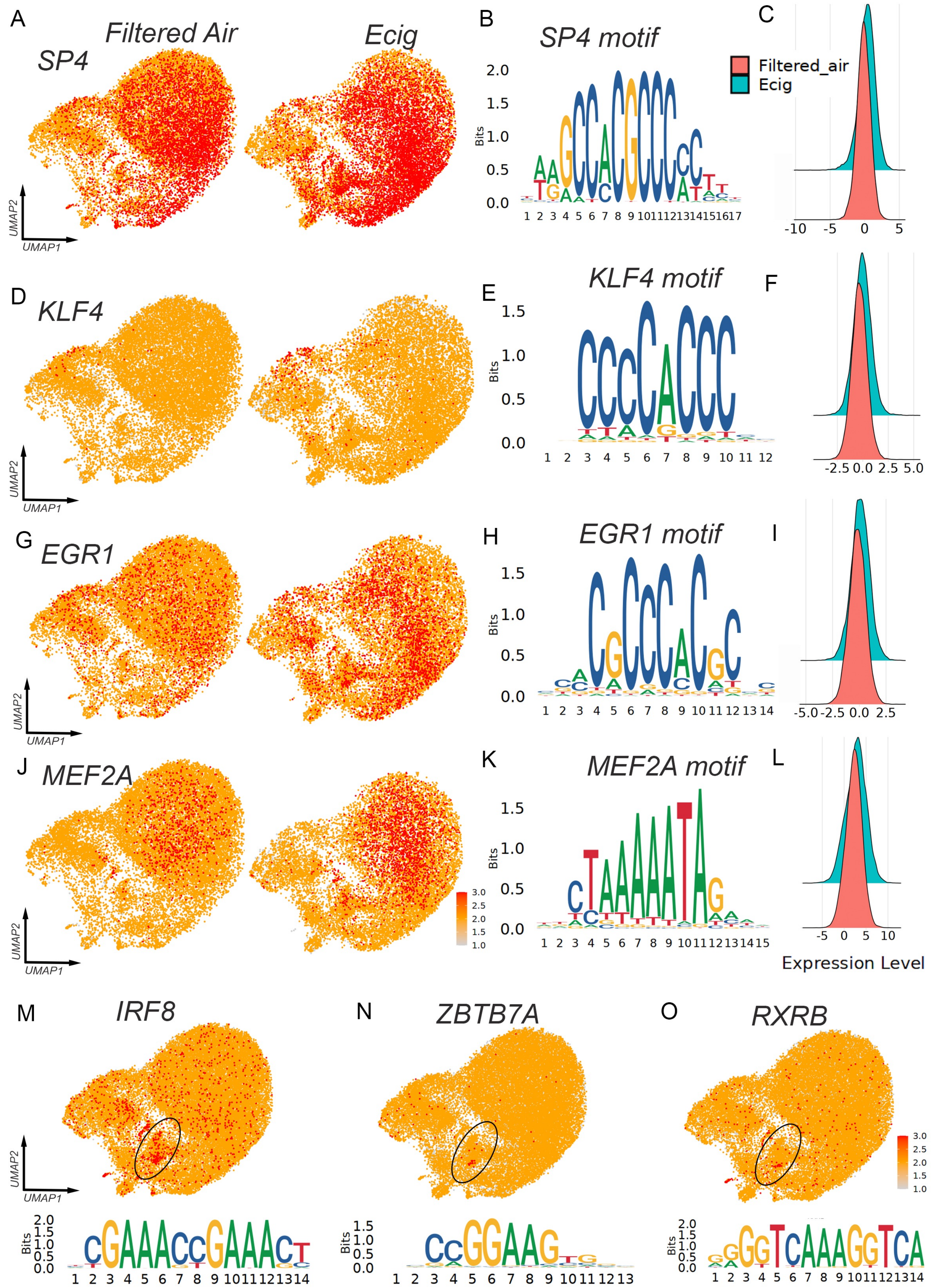

### Supplemental Figure 7

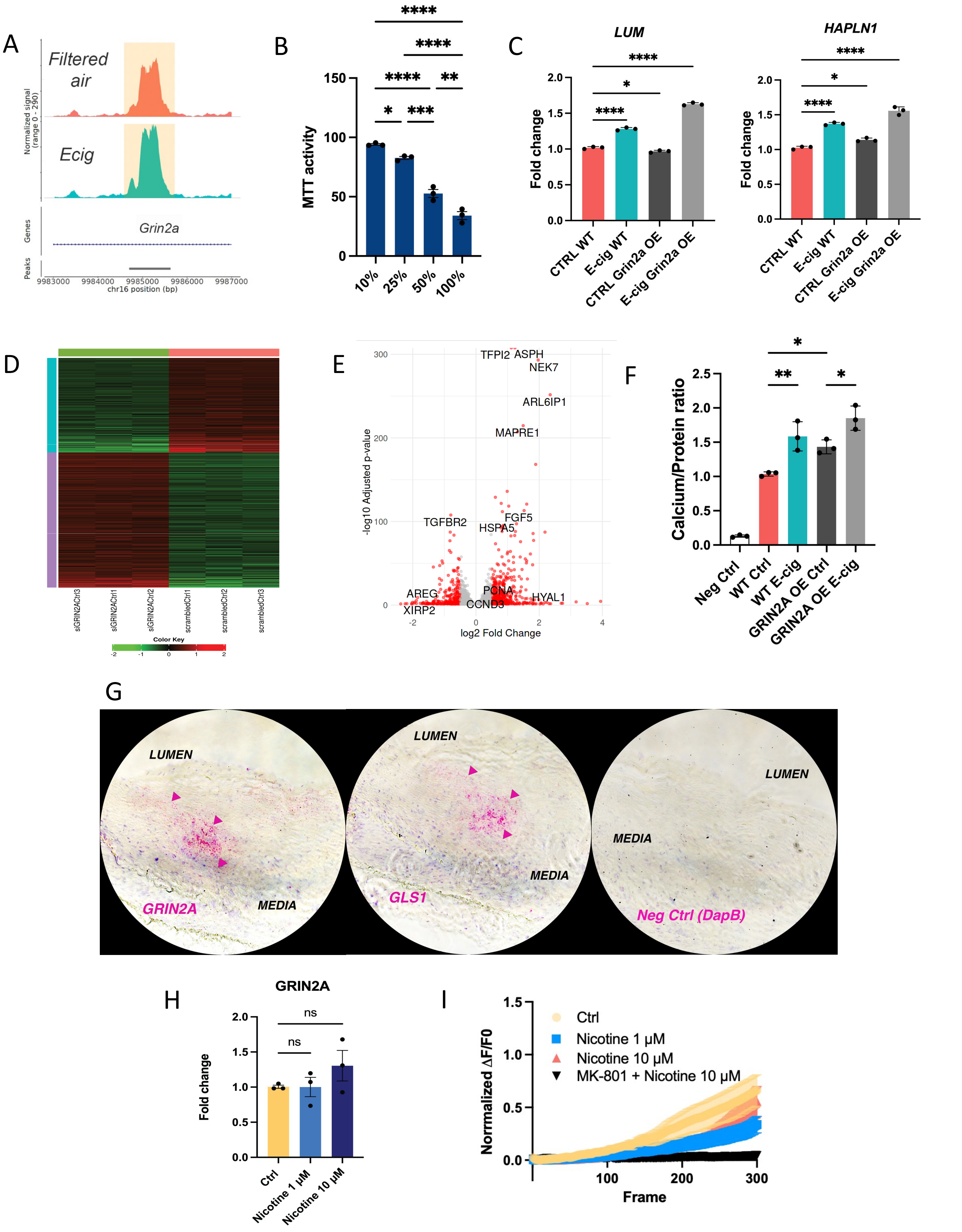
