## Supplemental Figure Legends for "Chronic Electronic Cigarette Exposure Promotes Atherosclerosis and Chondrogenic Modulation of Smooth Muscle Cells"

**Supplemental Figure 1. E-cigarette exposure does not impact systemic lipid levels.** (A) Urine 3-OH cotinine concentrations after 16-week inhalation of E-cig aerosol, P<0.0001. Mice were sacrificed 24h following the last exposure. (B) Total cholesterol was assessed in plasma samples, P=0. 84.Triglyceride levels were assessed in plasma samples, P=0.74. LDL levels were assessed in plasma samples, P=0.80. HDL levels were assessed in plasma samples, P=0.88. (C) Quantification of Oil Red O positive area. (D) Representative aortic root sections from Control and E-cig exposed mice stained for Oil Red O. Scale bar: 250µm.Data in (A,B,C) were analyzed by two-tailed T-test. Data represent mean +/- SEM. (n = 8-12 mice per group).

**Supplemental Figure 2.** (A) UMAP of scRNAseq data with cell clusters split by exposure type. (B) Volcano plot showing differential gene expression analysis by cell type and exposure type for Fibroblasts (Fibro), Macrophage (Mac), and Endothelial cells (Endo). (C) GO-term enrichment analysis for differentially expressed genes by exposure for SMC subset; Parentheses reflect distinct genes included in pathway enrichment analysis. (D) Dotplot gene expression for top marker genes for SMC clusters. (E) Representative images of RNA scope for *Fbln1 and Cxcl12* and *C3* in atherosclerotic plaque of the aortic root in 16-week E-cig exposed mice. The images are representative of three experiments, and the scale bars represent 50µm. Blue arrows highlight cells at the lesion and the media. (F) SMC-monocyte co-culture assay showing relative Thp1 monocyte migration across treatment groups. Data are expressed as mean +/- SEM. With p-value obtained using one-way ANOVA with Tukey's multiple comparisons post-hoc test, (no HCASMC vs Ctrl P= P<0.0001, Ctrl vs E-cig P=0.0002, Ctrl vs *C3*KD P=0.079, *C3*KD vs *C3*KD E-cig P=0.98, E-cig vs *C3*KD E-cig P<0.0001). Dots represent three technical replicates from three biologically independent samples.

**Supplemental Figure 3. Peak to gene relationship and biological effect highlights unfolded protein and cellular response to stress pathways with E-cig exposure.** (A) Volcano plot of differentially accessible peaks by E-cig exposure within Smooth muscle cells (SMC), Fibroblasts (Fibro), Endothelial cells (Endo) and Macrophage (Mac). (B) GO molecular function and (C) GO biological process of activated enhancer peak genomic region to gene relationship analysis by GREAT for E-cig exposure.

**Supplemental Figure 4.** GO biological processes for differentially accessible peaks using GREAT for (A) macrophage cells, (B) Fibroblasts, and (C) Endothelial cells.

**Supplemental Figure 5.** (A) UMAPs of scATACseq data from atherosclerotic aortic root and ascending aorta at 12 weeks of high fat diet with E-cigarette exposure in SMC lineage tracing mice (SMC-LnT: *Myh11*$^{CreERT2}$, *ROSA*$^{tdTomato}$, *ApoE*$^{-/-}$) grouped by cluster in all cells and in SMC subset grouped by cluster. Feature plot of *Myh11* gene activity in SMC subset. (B) EnrichR KEGG Pathway analysis of top marker genes by gene activity and GREAT analysis by chromatin activity for cluster '2' in scATACseq SMC subset. (C) EnrichR KEGG Pathway and Reactome analysis of top marker genes by gene activity for cluster "5" in scATACseq SMC subset. (D) EnrichR KEGG Pathway and Reactome analysis of top marker genes by gene activity for cluster "7" in scATACseq SMC subset

**Supplemental Figure 6. E-cigarette enhances transcription factor motif accessibility in vascular SMC.** Featureplots of scATACseq SMC subset for motif accessibility split by exposure, motif sequence, and ridge plot for motif accessibility split by exposure for SP4 (A-C), KLF4 (D-F), EGR1 (G-I), and MEF2A (J-L). Featureplots of scATACseq SMC subset for *'nicotine'* specific cluster markers and sequence for (M) IRF8, (N) ZBTB7A, and (O) RXRB.

**Supplemental Figure 7. GRIN2A regulates transcriptional reprogramming and glutamatergic pathway activation in response to E-cigarette exposure in HCASMCs and human atherosclerosis.** (A) Coverage plot of representative glutamate receptor gene *Grin2a* region split by exposure in SMC subset. (B) HCASMCs were treated with E-cig extract (10%, 25%, 50% or 100%) for 24 h, and cytotoxicity was determined by MTT assay. Data are expressed as mean +/- SEM. With p-value obtained using one-way ANOVA with Tukey's multiple comparisons post-hoc test, (10% vs 25% P=0.498, 10% vs 50% P<0.0001, 10% vs 100% P<0.0001, 25% vs 50% P=0.0003, 25% vs 100% P<0.0001, 50% vs 100% P=0.0049). (C) Fibroblast (*LUM*) and chondrocyte (*HAPLN1*) markers were evaluated by quantitative RT-PCR with *GRIN2A* overexpression in HCASMC. Data were expressed as mean +/- SEM. With p-value obtained using one-way ANOVA with Tukey's multiple comparisons post-hoc test, For *LUM* (Ctrl WT vs E-cig WT P=0.0674, Ctrl WT vs Ctrl *GRIN2A* OE P=0.99, Ctrl WT vs E-cig *GRIN2A* OE P=0.0003), For *HAPLN1* (Ctrl WT vs E-cig WT P=0.0013, Ctrl WT vs Ctrl *GRIN2A* OE P=0.13, Ctrl WT vs E-cig *GRIN2A* OE P<0.0001).Dots represent three technical replicates from three biologically independent samples. (D) Heatmap displaying differentially expressed genes in comparing si*GRIN2A* Ctrl versus scrambled Ctrl samples from bulk RNA-sequencing data in HCASMCs. (E) Volcano plot of differentially expressed genes by *GRIN2A* KD, highlighting a few significantly regulated genes including *TFPI2, ASPH, HSPA5, XIRP2, and AREG*. (F) Rate of calcification with *GRIN2A* overexpression in HCASMCs grown in calcification media. Data were expressed as mean +/- SEM. With p-value obtained using one-way ANOVA with Tukey's multiple comparisons post-hoc test, (Ctrl WT vs E-cig WT P=0.0036, Ctrl WT vs Ctrl *GRIN2A* OE P=0.030, Ctrl WT vs E-cig *GRIN2A* OE P=0.002). Dots represent three technical replicates from three biologically independent samples (G) RNA scope in situ hybridization of *GRIN2A (left), GLS1 (center), negative control (bacterial DapB, right)* in coronary artery sections from heart transplant recipients with atherosclerosis. The image is representative of five experiments in serial sections, and the scale bar represents 50μm. Red arrows highlight RNA expression in the lesion. (H) Expression of *GRIN2A* was evaluated in HCASMC by quantitative RT-PCR following nicotine treatment. Data were normalized relative to Ctrl and expressed as mean +/- SEM. With p-value obtained using one-way ANOVA with Dunnett's multiple comparisons post-hoc test, (Ctrl vs Nicotine 1μM P=0.99, Ctrl vs Nicotine 10μM P=0.33). Dots represent three technical replicates from three biologically independent samples. (I) Real-time calcium influx in HCASMCs measured using the FLUO-4 AM probe following treatment with 1μM and 10μM Nicotine or MK-801 plus 10μM nicotine. Calcium dynamics were recorded immediately after treatment for 300 seconds using an ECHO Revolve microscope. Data are expressed as ΔF/F0 where F0 is baseline fluorescence intensity of the first 10 frames and ΔF is the change in mean fluorescence intensity.

**Supplemental Table**

**Table 1. Clinical characteristics of the patients included in this study**

**Table 2. Differentially regulated genes (adjP <0.05) following *GRIN2A* siRNA knockdown in HCASMC**
