## Supplemental Materials and Methods for "Chronic Electronic Cigarette Exposure Promotes Atherosclerosis and Chondrogenic Modulation of Smooth Muscle Cells"

**Expanded methods:**

**Mouse strain -** The animal study protocol was approved by the Administrative Panel on Laboratory Animal Care (APLAC) at Stanford University. SMC-specific lineage tracing was generated by a well-characterized BAC transgene that expresses a tamoxifen-inducible Cre recombinase driven by the SMC-specific *Myh11* promoter ($Tg^{Myh11\text{-}CreERT2}$; 019079; Jackson Laboratory). These mice were bred with a floxed-stop-flox *tdTomato* fluorescent reporter line (B6.Cg-*Gt(ROSA)26Sor*$^{tm14(CAGtdTomato)Hze}$/J; 007914; Jackson Laboratory) to allow SMC-specific lineage tracing to generate the SMCLnT/WT mice. All mice were bred onto the C56BL/6, ApoE-/- background. As the Cre-expressing BAC was integrated into the Y chromosome, all lineage tracing mice in the study were male.

We conducted all animal experiments in accordance with institutional guidelines and report our in vivo studies in compliance with the ARRIVE reporting recommendations. For each experiment, the species, strain, age, sex, and source of animals are specified in the Methods. Animals were randomized to exposure/treatment groups using a prespecified allocation scheme, with allocation concealment where feasible, and investigators were blinded to group assignment during outcome assessment and data analysis when applicable. Prespecified inclusion and exclusion criteria (defined a priori) were applied consistently, and all excluded animals (including exclusions occurring after randomization) are reported with reasons. Statistical methods, including the unit of analysis, sample sizes, and criteria for significance, are described for each experiment.

**Induction of lineage marker by Cre recombinase -** Two doses of tamoxifen (Sigma-Aldrich, #T5648), at 0.2 mg $g^{-1}$ of body weight, were administered by oral gavage at 8 weeks of age, with each dose separated by 48 hours. Approximately 48 hours after the second dose of tamoxifen, high fat diet (HFD) was started (Dyets, #101511, 21% anhydrous milk fat, 19% casein, 0.15% cholesterol).

***Whole body E-cig aerosol exposure* -** A whole-body chamber of the inExpose™ system (Scientific Respiratory Equipment Inc. [SCIREQ], Montreal, QC, Canada) was used to deliver E-cig aerosol. Mice were randomly assigned to filtered air-exposed controls or Juul Virginia Tobacco-exposed groups. Juul Virginia Tobacco pods (5.0% nicotine by weight, 59 mg/mL) were selected due to their high nicotine content and widespread use among U.S. adolescents and young adults, ensuring translational relevance. Starting at 8 weeks of age (following tamoxifen administration), mice were exposed to aerosol for 2 hours per day, 3 days per week, for 12 or 16 weeks. Control mice were housed in individually ventilated cages equipped with HEPA-grade filters and were not placed in the exposure chamber to minimize potential confounding from residual aerosol contamination within the chamber system.
Aerosol was generated at a frequency of 1 puff per minute and duration of 3.3-4 seconds per puff.[1] The mice received 120 puffs/day over 2 hours, comparable to the average puff count of a human E-cigarette user.[2]

**Nicotine and Aerosol Chemical Characterization -** Nicotine in the aerosol phase, along with other chemical components (propylene glycol and aldehydes including acrolein, formaldehyde, and acetaldehyde) were quantified using selective ion flow tube mass spectrometry (SIFT-MS;

Syft Technologies) as previously described.[1] For each pod, six 70 mL puffs were collected into 0.5 L Tedlar bags, resulting in an 80–90% full bag capacity. Room air-filled bags served as controls. For each analyte, concentrations (in ppb) were averaged across 15 ms timeframes. Measured nicotine concentrations in the Tedlar bags ranged from 700 to 2000 ppb per 6 puffs, corresponding to an approximate per-puff concentration of 117 to 333 ppb.

**Gravimetric Determination of Total Particulate Matter (TPM) -** Total particulate matter (TPM) refers to the mass of particulate-phase aerosol components collected from the aerosol stream during puffing experiments. TPM was determined gravimetrically by weighing filter pads before and after exposure on an analytical scale. The average mass increase across filters was 150 mg per pod. TPM per puff was calculated as 150 mg total TPM divided by 120 puffs, yielding an average of 1.25mg/puff.

**Single-cell capture and library preparation and sequencing -** Immediately after sacrifice, mice were perfused with PBS. The aortic root was excised, and tissue were washed three times in PBS, placed into an enzymatic dissociation cocktail (2 U $ml^{-1}$, Liberase (Sigma-Aldrich, #5401127001) and 2 U $ml^{-1}$ elastase (Worthington, #LS002279) in Hank's Balanced Salt Solution (HBSS) and minced. After incubation at 37 °C for 1 hour, the cell suspension was strained and then pelleted by centrifugation at 500*g* for 5 minutes. The enzyme solution was then discarded, and cells were resuspended in fresh HBSS. To increase biological replication, multiple mice were used to obtain single-cell suspensions at each time point. For each scRNA capture, two mice were used. Cells were FACS sorted based on tdTomato expression. tdT-positive cells (considered to be of SMC lineage) and tdT-negative cells were then captured on separate but parallel runs of the same scRNA-Seq workflow and datasets were later combined for all subsequent analyses. For single cell ATAC, tdT- positive cells and tdT-negative cells were pooled at a 1:1 ratio, collected in BSA-coated tubes, and nuclei isolated per 10X recommended protocol, and captured on the 10X scATAC platform. Single-cell suspensions were loaded onto 10x GEM X/H chips and libraries were prepared using the Chromium Single Cell 3' RNA V4 and Chromium Single Cell ATAC V2 kits. Libraries were sequenced on an Illumina NovaSeq6000 platform, targeting a sequencing depth of 40,000–50,000 reads per cell for scRNA-seq and 75,000 reads per cell for scATAC-seq.

**Analysis of Single-Cell Sequencing Data *scRNAseq* -** Fastq files from each exposure group capture for tdT-positive (6 total tdT-positive libraries, 3 FA control, 3 E-cig) and tdT-negative (5 total tdT-negative libraries, 3 FA, 2 E-cig) cells were aligned to the reference genome (mm10) individually using CellRanger Software (10x Genomics). Dataset was then analyzed, and captures were integrated using the R package Seurat[3]. The dataset was trimmed of cells with an nFeature_RNA could of less then 2000, and genes expressed in fewer than 50 cells. The number of genes, number of unique molecular identifiers and percentage of mitochondrial genes were examined to identify outliers. As an unusually high number of genes can result from a 'doublet' event, in which two different cell types are captured together with the same barcoded bead, cells with >7500 genes were discarded. Cells containing >7.5% mitochondrial genes were presumed to be of poor quality and were also discarded. The gene expression values then underwent library-size normalization and normalized using established Single-Cell Transform function in Seurat. Principal component analysis was used for dimensionality reduction, followed by clustering in principal component analysis space using a graph-based clustering approach via Louvain algorithm. No batch correction was needed for analysis. UMAP was then used for two-dimensional visualization of the resulting clusters. Analysis, visualization and quantification of gene expression and generation of gene module scores were performed using Seurat's built-in function such as "FeaturePlot", "VlnPlot", and "FindMarker."

***scATACseq* -** Fastq files from each exposure group with tdT-positive and tdT-negative cells pooled in a 1:1 mixture (total 5 libraries, 2 FA control, 3 E-cig) were aligned to the reference ATAC genome (mm10) individually using CellRanger Software (10x Genomics). Individual datasets were aggregated and peak calling was performed using the CellRanger aggr command without subsampling normalization. The aggregated dataset was then analyzed using the R package Signac[3]. The dataset was first trimmed of cells containing fewer than 1000 peaks, and peaks found in fewer than 10 cells. The subsequent cells were then again filtered based off TSS enrichment, nucleosome signal, and percent of reads that lies within peaks found within the larger dataset. Cells with greater than 100,000 reads within peaks or fewer than 1000 peaks, <2 transcription start site (TSS) enrichment, or <25% reads within peaks were removed as they are likely poor-quality nuclei. The remaining cells were then processed using RunTFIDF(), RunSVD() functions from Signac to allow for latent semantic indexing (LSI) of the peaks[4,5], which was then used to create UMAPs. No batch correction was required for analysis. Differentially accessible peaks between different populations of cells were found using FindMarker function with default Wilcoxon Rank Sum testing with a reduced pseudocount.use value of 0.1, using number of peaks as latent variable to correct for depth. Motif matrix was obtained from JASPAR 2020, aligned onto BSgenome.Mmusculus.UCSC.mm10. Accessibility analysis around transcription factor motifs was performed using ChromVar[6]. Merging of scRNA and ATAC data was performed using Pseudo-expression of each gene created using scATACseq GeneActivity function to assign peaks to nearest genes expressed in the scRNA dataset, and mapped onto each other using Canonical Correlation Analysis. Label transfer was then performed following standard protocol.

**Pathway analyses of differential genes and peaks -** We performed pathway enrichment analyses on both differentially expressed genes (DEGs) and differentially accessible chromatin regions. For DEGs, we used Enrichr to identify enriched Gene Ontology (GO) terms, pathways (KEGG, Reactome), and transcription factor targets[7]. Significantly enriched pathways were filtered based on adjusted $p < 0.05$. For differentially accessible peaks derived from scATAC-seq, we utilized the Genomic Regions Enrichment of Annotations Tool (GREAT) version 3.0 to infer putative biological functions based on peak-to-gene annotations and regulatory domains[8]. We used the default "Basal plus extension" setting, which assigns regulatory domains to genes based on 5 kb upstream and 1 kb downstream of the transcription start site, plus distal regions extending up to 1 Mb, excluding overlaps with neighboring genes.

**Preparation of mouse aortic root sections -** Immediately after sacrifice, mice were perfused with 4% PFA. The mouse aortic root was excised and immersed in 4% PFA at 4C overnight.

After passing through a sucrose gradient, tissue was frozen in OCT (Thermo Fisher Scientific, #4585) to make blocks. Blocks were cut into 7um-thick sections for further analysis.

**Immunohistochemistry (IHC) -** Slides were prepared and processed as described previously[9]. Sections were then incubated overnight at 4°C with anti-SM22alpha rabbit polyclonal primary antibody (Abcam, #ab14106, 1:300 dilution), or anti-CD68 rabbit polyclonal antibody (Abcam, #ab125212, 1:300 dilution). Sections were washed for 5 minutes x2 with TBS and then incubated with the Rabbit-on-Rodent HRP Polymer (Biocare Medical, #RMR622) for 30 minutes at room temperature (RT). Sections were washed x2 with TBS and then incubated with the Betazoid DAB chromogen reagents (Biocare Medical, #BDB2004) for 4 minutes at RT. Sections were washed x2 in DI water and air-dried, followed by mounting with Eco mount (Biocare Medical). Oil Red O (ORO) staining (Sigma-Aldrich, #01391) was conducted to assess lipid content. For the alkaline phosphatase enzymatic assay, Ferangi Blue Chromogen kit (Biocare Medical, #FB813H) was used as instructed. Processed sections for alkaline phosphatase staining and ORO staining were visualized using ECHO Revolve microscope while IHC sections were visualized with a Leica DM5500B microscope. Areas of interest were quantified using ImageJ (NIH) software and compared using a two-sided t-test. The lesion cap was defined as 30µm segment from the luminal surface as previously described[9].

**RNAscope -** Slides were processed according to the manufacturer's instructions, and all reagents were obtained from ACD Bio. Slides were washed once in PBS, then immersed in 1Å~ Target Retrieval reagent at 98°C for 6 min. Slides were washed twice in deionized water, immersed in 100% ethanol and air dried, and sections were encircled with a liquid-blocking pen. Sections were incubated with Protease Plus reagent for 30 min at 40°C, then washed twice with deionized water. Sections were incubated with probes against mouse Grin2a, Fbln1, Cxcl12 and *C3* or a negative control probe or against human GRIN2A and GLS for 4h at 40 °C. RNAscope 2.5 HD assay- RED (ACD Bio, #322350) was performed per the manufacturer's instructions.

**Culture of human coronary artery smooth muscle cell (HCASMC) -** Primary human coronary artery smooth muscle cells (HCASMC) were purchased from Cell Applications, Inc. (#350-05a) and were cultured in complete smooth muscle basal media (Lonza, #CC-3182 and #CC-4149) according to the manufacturer's instructions. All experiments were performed with HCASMC between passages 5–8.

**Culture of immortalized HCASMC and THP1 Cell -** The immortalized hTERT HCASMC were obtained from Clint Miller lab. In general, the HCASMC were purchased from Cell Applications, Inc. (#350-05a) and immortalized by lentiviral transduction of the hTERT-IRES-hygro construct (Addgene Plasmid, #85140) in 10 µg/ml polybrene. Transduced cells were selected using 400 µg/ml of hygromycin (Gibco, #10687010) and maintained in SmBm medium for cell culture. The THP-1 cell line was purchased from company. THP-1 cells were cultured in RPMI 1640 medium (Corning, #15040CV) supplemented with 10% FBS (Sigma-Aldrich, #12306C).

**Knockdown and overexpression -** For the siRNA transfection, cells were grown to 70% confluence, then treated with siRNA or scramble control to final concentration of 10nM with Lipofectamine RNAiMax reagent (Thermo Fisher Scientific, #13778150) for 24 hours. The siRNA for GRIN2A was purchased from Origene (#SR301955). Cells were allowed to recover in SMC growth medium for 24 hours before cell harvest. For overexpression studies, the human *GRIN2A* overexpression construct was designed and synthesized by VectorBuilder (Vector ID: VB240131-1404auv). Lentiviral particles were generated in HEK293T cells via transient transfection using a second-generation packaging system. Cells were transfected with a 1:0.25:1 molar ratio of the packaging plasmid pCMV-dR8.91, the envelope plasmid pMD2.G, and the *GRIN2A* transfer plasmid using Lipofectamine 3000 (Thermo Fisher Scientific, #L3000015) in Opti-MEM (Gibco, #22600050), according to the manufacturer's instructions. Viral supernatants were collected 48 hours post-transfection, filtered through a 0.45µm filter, and used immediately for transduction. HCASMCs were seeded to reach ~60% confluence and transduced with viral supernatant for 6 hours in the presence of polybrene (8µg/mL). Following transduction, the medium was replaced with fresh growth media, and cells were incubated overnight prior to collection for downstream applications. For transwell migration assay, the siRNA for C3 was purchased from Horizon Discovery (NM_000064).

**RNA isolation and qRT-PCR -** RNA was isolated using RNeasy mini kit (Qiagen, #74104) and total cDNA was prepared using High-capacity RNA-to-cDNA kit (Applied Biosystems, #43-874-06). Gene expression levels were measured using Taqman probes (Invitrogen) for GRIN2A (*Hs00168216_m1*), *LUM* (*Hs00929860_m1*), *HAPLN1 (Hs01091999_m1)*, *GLS* (*Hs01014020_m1*), *GLUL*(*Hs00365928_g1*), *SLC1A3* (*Hs00904823_g1*), *SLC7A11 (Hs00921938_m1)* and quantified on a ViiA7 Real-Time PCR system (Applied Biosystems, Foster City, CA) and normalized to *GAPDH* levels.

**RNA-sequencing -** Three replicates were used for each sample. The RNA was processed and analyzed as described previously. Briefly, RNAs were sent to Novogene for sample QC, library preparation, and sequencing. All samples passed QC, and 250– 300 bp insert cDNA libraries were prepared for each sample. Subsequently, sequencing was performed on a NovaSeq X Plus platform with 20M paired reads. DESeq2 was used for differential expression analysis using the Wald test. Additionally, the web-based tools iDEP2.0(http://bioinformatics.sdstate.edu/idep/) was used for analysis of the RNA-Seq data and visualization using the counts data generated from Feature Counts.

***In-vitro HCASMC aerosol exposure to E-cig* -** The inExpose™ inhalation exposure system with the Flexiware software (v8.3) (SCIREQ, Montreal, QB, Canada) was used for generating E-cig aerosol extract for exposure to HCASMC. In brief, *Juul* pod containing 5.0% (59 mg/ml) nicotine by weight and a Virginia tobacco flavor was purchased from the *Juul* online store. *Juul* device was attached to the JUUL E-cig compatible extension device provided with the inExpose equipment. The E-cig aerosol was drawn into a small sterile glass bottle, containing 20 mL of appropriate sterile complete cell culture medium with serum. 70 puffs were vaporized into the medium according to the following parameters: one 91ml puff of 4-second duration every 60-seconds[10,11]. Flow rates were monitored and regulated via an integral flow meter (Key Instruments, Trevose, PA, USA). The E-cig vapor extract was collected, and cell viability was assessed at different concentrations. For in vitro experiments, aerosol extracts were diluted to 25% (v/v), corresponding to a final nicotine concentration of approximately 37-50 µM.

**Cytotoxicity Assessment -** HCASMC were seeded into 24-well plates at a density of 50,000 cells/well. To assess the effect of E-cig extract on cytotoxicity, a 3-(4,5-dimethylthiazol-2-yl)-2,5-diphenyltetrazolium bromide (MTT) assay was used to measure the relative amount of metabolically active cells. Cells were exposed to undiluted E-cig aerosol extract (100%) and to serial dilutions (50%, 25%, 10%). A MTT assay kit was then used according to the manufacturer's specifications (Abcam, #ab211091). Following the MTT reaction, absorbance was measured at a wavelength of 570nm using a SpectraMax iD3 series Microplate Reader (Molecular Devices, San Jose, CA, USA). Subsequently, the viability of the E-cig-treated cells was expressed as a percent of untreated control cells.

**Nicotine quantification in E-cig aerosol extracts.** Nicotine concentrations in E-cig aerosol extracts used for in vitro experiments were quantified by LC–MS/MS using a Sciex 7500+ triple quadrupole mass spectrometer with a phenyl-hexyl analytical column and a water/methanol solvent system. Calibration standards were processed in parallel with experimental samples, and measurements were performed within the linear dynamic range of the assay. Nicotine concentrations measured in aerosol extracts ranged from 147 to 202 μM across independent preparation batches (mean ± SD: 174.5 ± 27.2 μM).

**HCASMC Phenotypic Assays -** For the proliferation assay, The HCASMC were treated with siRNA for GRIN2A as described in "Knockdown and overexpression" and then exposed to E-cig aerosol extract for 24h. EdU (10μM) was introduced into the cell culture 2 hours before assay for uptake. The protocol for Click-iT Plus EdU proliferation kit (Invitrogen, #C10637,) was followed as instructed. To assay for calcification, HCASMC was exposed to calcification media with 10mM beta-glycerophophate (Sigma-Aldrich, #G9422), 100nM Insulin (Roche, #11376497001), 50ug/ml ascorbic acid (Thermo Fisher Scientific, #036237) and 8mM $CaCl_2$ (Thermo Fisher Scientific, #J63122). The HCASMC were treated with siRNA and lentivirus for GRIN2A as described in "Knockdown and overexpression" and then exposed to calcification media with 0.4% FBS for 6 days. The calcified cells were treated with 0.6N hydrochloric acid for 24h, then the supernatant was collected for calcium assay, and the cell layer was collected for protein quantification. The Calcium Colorimetric Assay Kit (Abcam, #ab102505) was used for quantification of calcium, then normalized to total protein content. Glutamate release from HCASMCs was measured using the Amplex Red Glutamic Acid/Glutamate Oxidase Assay Kit (Invitrogen, #A12221). HCASMCs were cultured in vascular cell basal medium (ATCC, #PCS-100-030) without phenol red, supplemented with 0.1% fatty acid–free BSA (Sigma-Aldrich, #A3311), and maintained under low-serum conditions (1.25% FBS, final concentration) to minimize background glutamate. Cells were treated with 25% E-cig aerosol extract or vehicle control for 24h. Following treatment, culture supernatants were collected, clarified by centrifugation, and analyzed according to the manufacturer's instructions. Fluorescence was measured at 590 nm using a SpectraMax iD3 microplate reader (Molecular Devices). Calcium flux imaging was conducted by incubating HCASMCs with the Fluo-4 AM calcium probe (Thermo Fisher Scientific, #F14201) for 30 minutes at 37°C. Following incubation, cells were washed and equilibrated in $Mg^{2+}$-free HEPES-buffered saline solution. This solution was specifically formulated to exclude glutamate to avoid confounding activation of glutamate receptors and to facilitate NMDAR activation by relieving voltage-dependent $Mg^{2+}$ blockade at resting membrane potential. The buffer consisted of (in mM): 145 NaCl, 5 KCl, 2 $CaCl_2$, 10 HEPES, 10 glucose; pH adjusted to 7.4 with NaOH. Cells were then treated with one of the following conditions: 25% E-cig aerosol extract, 10μM MK-801 (Sigma-Aldrich, #M107) combined with E-cigarette extract, 100μM NMDA (Tocris, #0114) plus 30μM D-serine (Cayman Chemical, #31197) or Nicotine (Sigma-Aldrich, #N0267). Calcium dynamics were recorded immediately after treatment for 300 seconds using an ECHO Revolve microscope (ECHO BICO Company, San Diego, CA, USA). For transwell migration assay, hTert-immortalized HCASMC were grown to 70% confluence in 24-well plate, then transfected with C3 siRNA or scramble control to final concentration of 20nM with RNAiMax (Invitrogen, #13778075) for 24 hours. Then change to SmBm medium for 24h, followed by E-cig aerosol extract treatment or SmBm control for 24h. Then a cell migration assay was performed to study the migration behavior of THP-1 macrophage cells. Cell culture inserts for 24-well plate containing a membrane with pores of 8μm (Corning, #353097) were used. Fifteen thousand THP-1 macrophages in 200μL of 1% FBS RPMI-1640 were loaded into both inserts and 24-well plates. Control cells were treated with the vehicle alone (DMSO). The transwell chambers were then incubated at 37°C in 5% $CO_2$ for 24h to evaluate cell migration. Then, the inserts were removed and THP-1 cells that migrated through the membrane to the lower chamber were quantified by imaging.

**Measurements of cotinine levels -** Following the 16 weeks exposure, 24h after the last exposure to the E-cigarette aerosol, mouse urine was collected via terminal bladder puncture and used for determination of cotinine levels using LC-LC/MS, performed by Creative Proteomics.

**Measurement of total cholesterol, triglycerides, LDL and HDL levels -** Total cholesterol, triglyceride, LDL and HDL were assessed in plasma samples obtained by cardiac puncture. All analysis was conducted by the Animal Diagnostic Laboratory at Stanford University.

**Human Samples -** Human coronary artery samples were obtained from the Stanford Department of Cardiothoracic Surgery Human Biorepository Tissue Bank from consenting patients, under protocol approval [Stanford IRB protocol number 67212]. All samples were from heart transplant recipients. Tissue procurement and use were conducted under protocols approved by the Stanford University Institutional Review Board. Patient identifiers, including Medical Record Numbers, were used only by the biobank for internal tracking and were not disclosed in the published data.
